# A Kiss-and-Ride Mechanism for Lipid Exchange at Membrane Contact Sites

**DOI:** 10.64898/2026.09.29.755298

**Authors:** Alena Balleková, Andrea Eisenreichová, Roman Holič, Peter Griač, Evžen Bouřa, Jana Humpolíčková

## Abstract

Lipid transfer proteins (LTPs) mediate non-vesicular lipid transport at membrane contact sites, yet how transient membrane tethering contributes to lipid transfer remains poorly understood. Here, we use the single-domain yeast phosphatidylserine transporter Osh6, a member of the oxysterol-binding protein–related protein family, to investigate tethering-dependent lipid transport. We show that Osh6 can access two transport modes depending on membrane composition: conventional shuttling between membranes and a tethering-dependent mode in which the opposing membrane facilitates cargo extraction from the donor membrane. Combining lipid transport and membrane-tethering measurements with kinetic analysis, lipid partitioning, and *in vivo* complementation, we provide evidence for a transient membrane-bridged intermediate during lipid transfer. Productive phosphatidylinositol 4-phosphate (PI4P) extraction destabilizes this bridged state, allowing cargo-loaded Osh6 to disengage and continue the transport cycle. We term this mechanism **kiss-and-ride**, in which transient membrane contact facilitates cargo extraction (“kiss”) before the cargo-loaded protein disengages (“ride”). Consistent with this model, PI4P transport is preserved in a lidless Osh6 mutant with impaired membrane dissociation, whereas phosphatidylserine transport is strongly reduced, revealing distinct mechanistic requirements for the two cargoes. Together, our findings reveal a tethering-dependent transport mode in a cup-like LTP and suggest that membrane contact sites can actively shape the mechanism of lipid transport rather than merely bringing donor and acceptor membranes into proximity.

## Introduction

Phospholipids play essential roles in membrane biogenesis and cellular signaling, and the specific lipid composition of each organelle strongly influences its function (1). Because most lipids are synthesized in the endoplasmic reticulum (ER), their distribution among cellular membranes must be tightly regulated (2–4). In addition to vesicular trafficking, lipids are transported between organelles by lipid transfer proteins (LTPs) (5–8).

LTPs primarily operate at membrane contact sites (MCSs), where two membranes are brought into close proximity by dedicated tethering proteins or by the lipid transporter itself (8, 9). Such proximity is thought to accelerate lipid exchange and contribute to transport directionality (10). Consistent with a functional role of membrane interactions, mutation of the distal membrane-binding site of the sterol transporter Osh4 compromises both protein function and yeast viability (11), while disruption of the interaction between the PS transporter Osh6 and its tethering partner Ist2 also affects yeast viability (12, 13).

Two structurally distinct mechanisms of non-vesicular lipid transport have been described (7, 14). Cup-like LTPs shuttle lipids within a hydrophobic binding pocket between donor and acceptor membranes. In contrast, rod-like proteins containing repeating β-groove (RBG) domains directly bridge membranes and form a continuous hydrophobic pathway for lipid movement (15). Whether cup-like LTPs can also exploit transient membrane bridging during lipid transfer remains unknown.

Here, we use the yeast phosphatidylserine (PS) transporter Osh6, a member of the oxysterol-binding protein–related protein (ORP) family (16, 17), to address this question. Osh6 is a well-characterized representative of cup-like LTPs, which transport lipids by enclosing individual cargo molecules within a hydrophobic binding pocket. Osh6 consists of a single lipid-transfer domain homologous to the transport domain of mammalian ORP5/8 (16, 18, 19). Osh6 transports PS from the ER to the plasma membrane (PM) in exchange for phosphatidylinositol 4-phosphate (PI4P) (20–22). PI4P is transported back to the ER and dephosphorylated by Sac1, while its PM pool is continuously replenished, providing the driving force for counter-gradient PS transport (23, 24).

PI4P binds Osh6 with higher affinity than PS (16, 25), allowing it to preferentially occupy the binding pocket after PS release and thereby suppress backward PS transport (21, 22). Membrane composition further regulates this exchange: PI4P is extracted more efficiently from highly negatively charged membranes, whereas PS extraction from such membranes is suppressed and PS release into charged membranes is favored (22).

Cargo-dependent changes in membrane affinity are a recurrent feature of LTPs (26, 27). In Osh6, the N-terminal lid acts as an electrostatic switch that regulates membrane association by shielding membrane-interacting surfaces upon cargo binding (28). Stepwise lid-shortening experiments show that the lid alters the electrostatic properties of Osh6 and impairs its dissociation from membranes. Because Osh6 can also tether membranes (11), lid shortening is expected to favor simultaneous interaction with two opposing membranes.

Here, we show that lid controls membrane tethering and differentially impacts PS and PI4P transport both *in vitro* and *in vivo*. Our results suggest that transient membrane tethering can participate directly in Osh6-mediated lipid transport, allowing Osh6 to engage donor and acceptor membranes simultaneously. Thus, in addition to canonical shuttling between membranes, Osh6 can access a membrane-bridged transport state. These findings reveal a tethering-dependent mode of lipid transport by a cup-like LTP and suggest that shuttle-based and membrane-bridged transport may represent complementary transport modalities within the ORP family.

## Results

The N-terminal lid of Osh6 controls both closure of the cargo-binding cavity and membrane dissociation (28). To investigate the relationship between lipid transport and membrane tethering, we prepared a series of progressively truncated Osh6 mutants: Osh6^Δ35^ lacks the disordered N-terminal segment, Osh6^Δ42^ additionally lacks the first N-terminal helix, and the Δ69 mutant lacks the entire lid (Osh6^lidless^). We compared their lipid extraction, transport, and membrane-tethering activities to determine how the lid affects the transport mechanism of Osh6.

To investigate how membrane composition and the presence of an opposing membrane affect Osh6-mediated lipid transport, we employed fluorescence cross-correlation spectroscopy (FCCS) (22, 29). We monitored either depletion of a cargo lipid (PS or PI4P) from donor LUVs (LUVs D) or its accumulation in acceptor LUVs (LUVs A). Donor or acceptor LUVs were labeled with DiD, while the cargo lipid was detected using fluorescent biosensors: C2_Lact_-CFP for PS (30, 31) and SidC-Atto488 for PI4P (32, 33). Biosensor binding to DiD-labeled cargo-containing LUVs results in correlated fluorescence fluctuations, quantified by the cross-correlation amplitude at zero lag time, *G*_cc_(0). Because biosensor responses depend on membrane composition, they were characterized separately for each lipid environment (34).

Experiments were performed in two complementary regimes. In the **extraction regime**, DiD-labeled donor LUVs were monitored, and cargo removal from the donor membrane resulted in a decrease in *G*_cc_(0). In the **transport regime**, DiD-labeled acceptor LUVs were monitored, and cargo accumulation in the acceptor membrane resulted in an increase in *G*_cc_(0). Thus, the transport regime directly reports lipid transfer from LUVs D to LUVs A, whereas the extraction regime reports only cargo removal from LUVs D and does not by itself reveal whether the extracted lipid remains bound to Osh6 or is subsequently transferred to LUVs A.

Under defined conditions, however, the extraction regime can also provide evidence for sustained lipid transport. The biosensor signal reports cargo depletion from donor LUVs. If this depletion exceeds the total binding capacity of Osh6 present in the assay, it requires repeated cycles of cargo extraction and release. At high initial cargo concentrations, however, the biosensor is saturated, and the first transport cycles therefore produce no detectable change in signal. Transport becomes detectable only after sufficient cargo depletion brings the biosensor into its responsive range.

### Acceptor-dependent PI4P removal from donor membranes

We first examined PI4P removal from donor membranes of different lipid compositions, including conditions resembling the large excess of PS over PI4P in the PM. Importantly, we distinguish between two modes of PI4P removal: **acceptor-independent removal**, in which Osh6 acquires PI4P from the donor membrane alone, and **acceptor-dependent removal**, in which efficient PI4P removal requires the presence of a second membrane. In the latter case, the acceptor membrane is not simply the destination for transported PI4P but contributes to its removal from the donor membrane.

We previously showed that, under conditions of excess PS in the donor membranes, Osh6 alone removes little PI4P from such membranes, whereas efficient PI4P transfer occurs when acceptor membranes are also present (22). We therefore asked whether and how the acceptor membrane facilitates PI4P removal from the donor membrane (Fig. **1A**).

**Figure 1.**
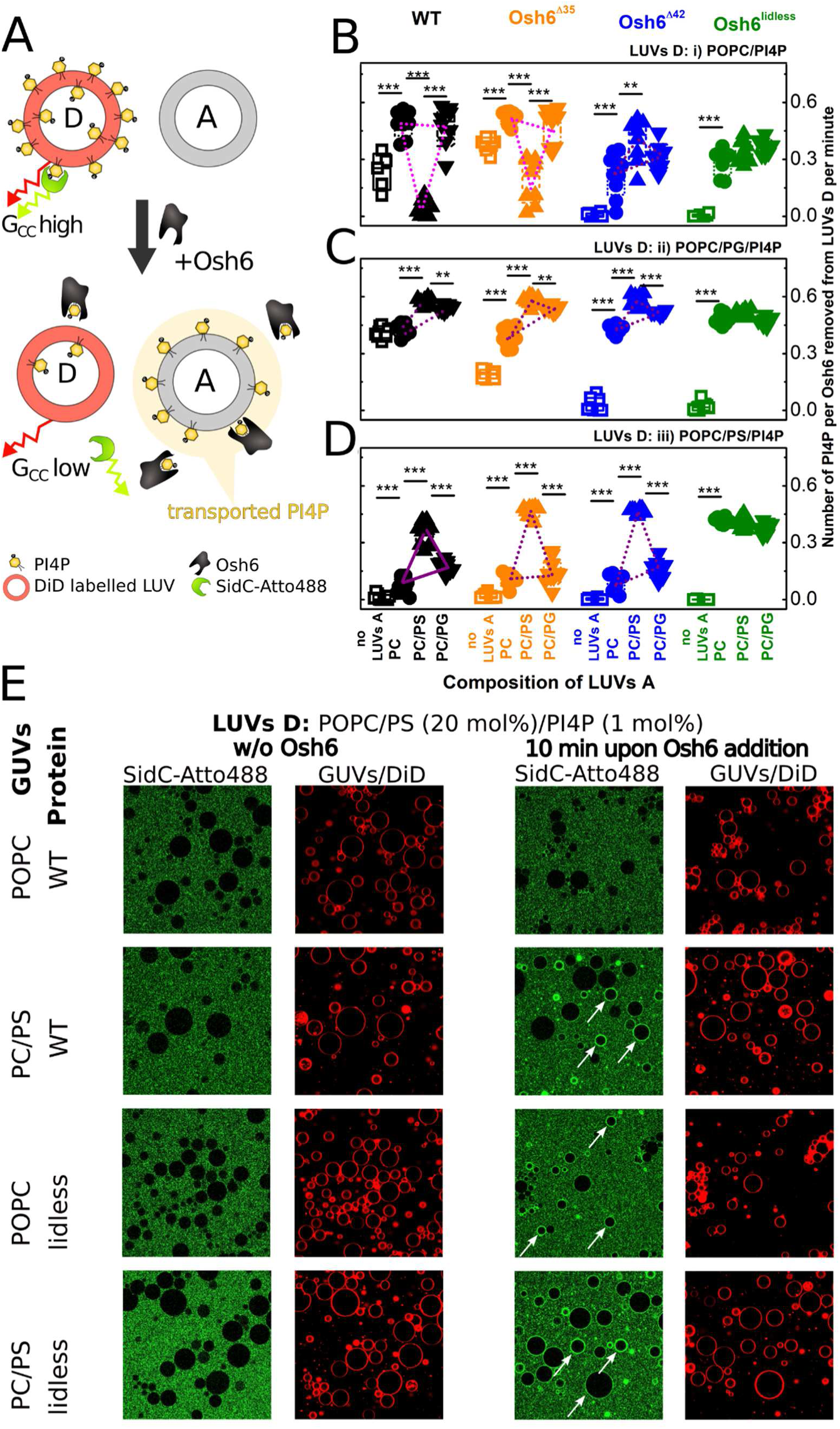
PI4P extraction and transfer by wild-type Osh6 and N-terminal truncation mutants. **A)** Scheme of the FCCS assay used to monitor PI4P removal from donor LUVs (LUVs D). LUVs D contain the fluorescent lipid tracer DiD, while PI4P is detected by the biosensor SidC-Atto488. Binding of SidC-Atto488 to PI4P-containing LUVs D produces correlated fluorescence fluctuations and a high cross-correlation amplitude, *G*_cc_(0). PI4P removal from LUVs D decreases SidC-Atto488 binding and consequently reduces *G*_cc_(0). **B–D)** Rates of PI4P removal from LUVs D in the absence of acceptor LUVs (LUVs A; open squares) or in the presence of LUVs A composed of POPC (circles), POPC/POPS (20 mol%; upward-pointing triangles), or POPC/POPG (20 mol%; downward-pointing triangles). Rates are expressed as PI4P molecules removed per Osh6 molecule per minute. Wild-type Osh6 and the Osh6^Δ35^, Osh6^Δ42^, and Osh6^lidless^ mutants are shown in black, orange, blue, and green, respectively. LUVs D were composed of POPC/PI4P (1 mol%) in **B**, POPC/POPG (20 mol%)/PI4P (1 mol%) in **C**, and POPC/POPS (20 mol%)/PI4P (1 mol%) in **D**. Solid and dotted violet lines and the magenta line highlight characteristic trends in PI4P removal across acceptor membrane compositions. Total lipid concentrations were 50 μM in LUVs D and 200 μM in LUVs A. PI4P concentration in LUVs D was 500 nM (250 nM in the outer leaflet accessible to Osh6); Osh6 and SidC-Atto488 concentrations were 250 nM and 80 nM, respectively. **E)** Visualization of PI4P transfer from donor LUVs composed of POPC/POPS (20 mol%)/PI4P (1 mol%) to GUVs composed of either POPC or POPC/POPS (20 mol%), mediated by wild-type Osh6 or Osh6^lidless^. PI4P transferred to GUVs was detected by recruitment of SidC-Atto488 (green), and GUV membranes were visualized with DiD (red). Total lipid concentrations were 250 μM in LUVs D and 12.5 μM in GUVs. PI4P concentration in LUVs D was 2.5 µM; Osh6 and SidC-Atto488 concentrations were 60 nM and 80 nM, respectively.

Three PI4P-containing donor LUVs (LUVs D) were used: (i) neutral POPC/PI4P (1 mol%) membranes (Fig. **1B**), (ii) negatively charged POPC/POPG (20 mol%)/PI4P (1 mol%) membranes (Fig. **1C**), and (iii) POPC/POPS (20 mol%)/PI4P (1 mol%) membranes containing an excess of the competing Osh6 cargo PS (Fig. **1D**).

Wild-type Osh6 readily removed PI4P from neutral and PG-containing donor membranes in the absence of acceptor LUVs (Fig. **1B,C**, black open squares). In contrast, little PI4P was removed from PS-containing donor membranes unless acceptor LUVs (LUVs A) were present (Fig. **1D**). The extent of PI4P removal then strongly depended on acceptor composition: neutral POPC LUVs A produced moderate removal, negatively charged POPC/POPG (20 mol%) LUVs A produced a stronger effect, and PS-containing POPC/POPS (20 mol%) LUVs A produced the strongest PI4P removal. This progressive increase is highlighted by the purple triangle in Fig. **1D**. Thus, under conditions of excess PS, both the presence and composition of the opposing membrane influence PI4P removal.

A similar, although weaker, acceptor dependence was observed with PG-containing donor membranes (Fig. **1C**), as highlighted by the dotted purple triangle. PI4P was removed even in the absence of LUVs A, but acceptor membranes further enhanced removal with a similar dependence on their composition. Thus, acceptor-independent and acceptor-dependent PI4P removal can operate simultaneously, with their relative contributions depending on the donor membrane environment.

Neutral donor membranes behaved differently (magenta wedge, Fig. **1B**). Here, PS-containing LUVs A inhibited PI4P removal, consistent with competition between PS and PI4P for the Osh6 binding pocket, whereas neutral and PG-containing LUVs A enhanced removal. Most notably, PS in the acceptor membrane had opposite effects depending on the donor: it inhibited PI4P removal from neutral donors but produced the strongest enhancement from negatively charged donors. Thus, the contribution of the acceptor membrane to PI4P removal depends strongly on the donor membrane environment.

Next, we used lid deletion mutants to dissect how the N-terminal lid affects this behavior. Osh6^Δ35^, missing only the N-terminal disordered region, closely resembled wild-type Osh6 under all conditions tested (Fig. **1B–D**, orange). Osh6^Δ42^, missing also the first helix of the lid, however, showed little PI4P removal from neutral or PG-containing donor membranes in the absence of an acceptor (Fig. **1B–D**, blue open squares). Addition of LUVs A restored PI4P removal and, across donor membrane compositions, produced the acceptor-composition dependence characteristic of the acceptor-dependent mode observed for wild-type Osh6 with negatively charged donors (purple triangles in Fig. **1B–D**).

The strongest acceptor dependence was observed for Osh6^lidless^ (Fig. **1B–D**, green). This mutant showed little PI4P removal from any donor membrane in the absence of LUVs A, but substantial removal when acceptor membranes were present. Unlike wild-type Osh6 and the shorter truncation mutants, however, PI4P removal by Osh6^lidless^ was largely independent of acceptor lipid composition. Because lid removal exposes a basic membrane-interacting surface and impairs membrane dissociation (22, 28), these results reveal a progressive shift from acceptor-independent toward acceptor-dependent PI4P removal as the N-terminal lid is shortened. In Osh6^lidless^, the presence of the second membrane itself, rather than its specific lipid composition, becomes the dominant factor enabling PI4P removal.

Because PI4P removal in the extraction regime does not by itself establish transfer to the acceptor membrane, we directly visualized PI4P accumulation in acceptor GUVs (Fig. **1E**). PI4P was transported from POPC/POPS (20 mol%)/PI4P (1 mol%) donor LUVs to either neutral POPC or PS-containing POPC/POPS (20 mol%) GUVs and detected by recruitment of SidC-Atto488 to the GUV membrane.

Consistent with the FCCS measurements, Osh6^lidless^ transferred PI4P to GUVs largely independently of their lipid composition, whereas wild-type Osh6 preferentially transferred PI4P to PS-containing GUVs. Thus, the acceptor-dependent PI4P removal observed by FCCS corresponds to lipid transfer to the acceptor membrane, while the strong dependence of this process on acceptor composition in wild-type Osh6 is largely lost in Osh6^lidless^.

### PS extraction and transport depend on membrane charge

Having identified acceptor-dependent PI4P removal, we next asked whether PS similarly depends on the presence of a second membrane. In cells, Osh6 transports PS from the weakly charged ER to the highly charged PM against its concentration gradient, while PI4P in the acceptor membrane promotes directionality by competitively suppressing backward PS transport (21, 22).

We first examined PS removal by wild-type Osh6 and the N-terminal truncation mutants from neutral and negatively charged donor membranes (Fig. **2A,B**). PS removal progressively decreased with N-terminal shortening and was consistently less efficient from negatively charged than from neutral membranes. No detectable PS removal was observed for Osh6^lidless^.

**Figure 2.**
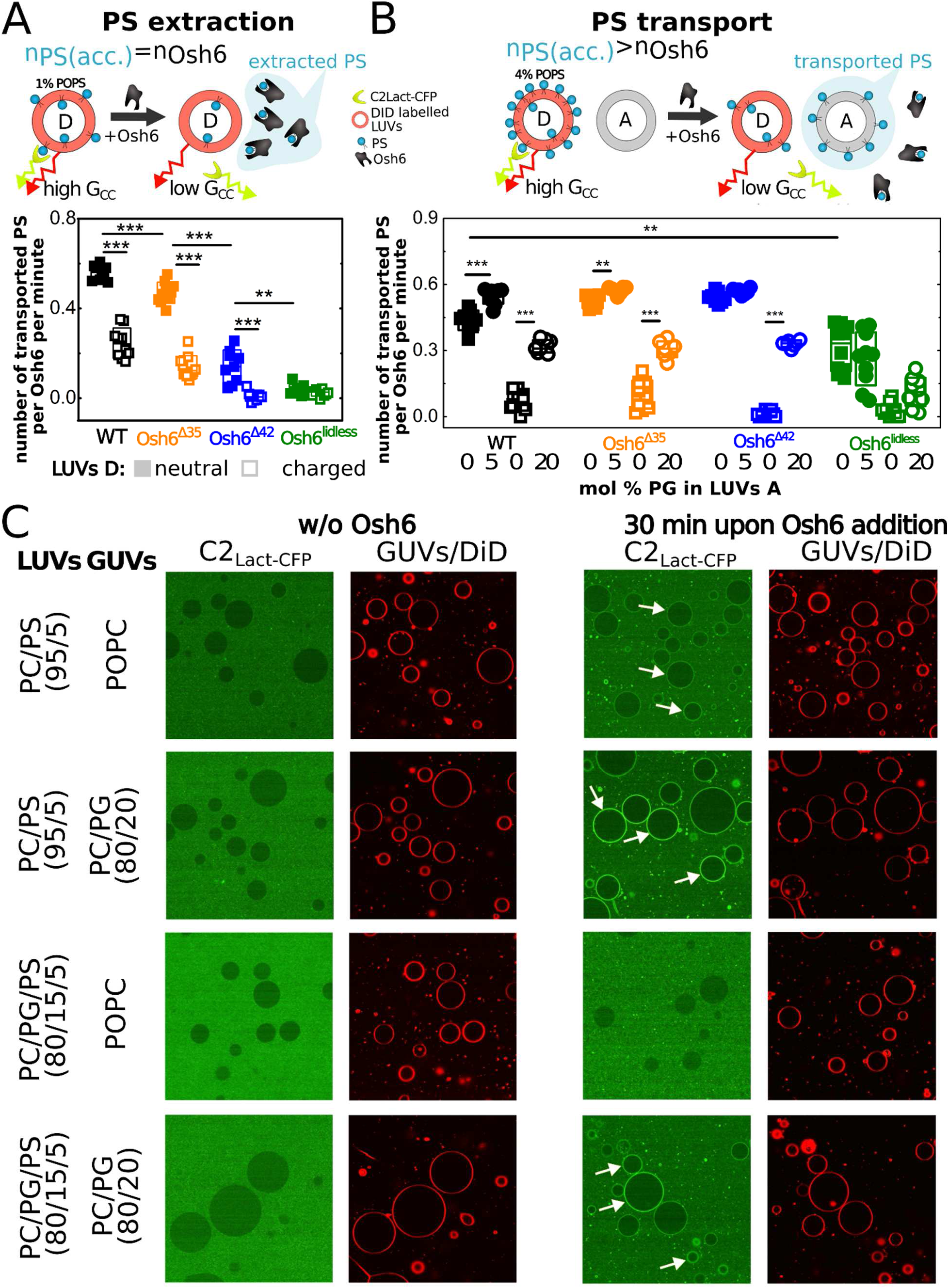
PS removal and transport by wild-type Osh6 and N-terminal truncation mutants. **A)** Scheme of the FCCS assay used to monitor PS removal from donor LUVs (LUVs D). LUVs D contain the fluorescent lipid tracer DiD, while PS is detected by the biosensor C2_Lact_-CFP. Binding of C2Lact-CFP to PS-containing LUVs D produces correlated fluorescence fluctuations. PS removal from LUVs D decreases C2_Lact_-CFP binding and consequently reduces the cross-correlation amplitude, *G*_cc_(0). **B)** Rates of PS removal from LUVs D composed of POPC/PS (1 mol%; neutral membranes, solid squares) or POPC/PG (20 mol%)/PS (1 mol%; negatively charged membranes, open squares). Rates are expressed as PS molecules removed per Osh6 molecule per minute. Wild-type Osh6 and the Osh6^Δ35^, Osh6^Δ42^, and Osh6^lidless^ mutants are shown in black, orange, blue, and green, respectively. Total lipid concentration in LUVs D was 50 μM. PS concentration in LUVs D was 500 nM (250 nM in the outer leaflet accessible to Osh6); Osh6 and C2_Lact_-CFP concentrations were 250 nM and 50 nM, respectively. **C)** PS transport rates (number of PS molecules transported to acceptor LUVs A per minute per Osh6 molecule). Donor LUVs D were composed of POPC/PS (4 mol%) (neutral membranes, solid symbols) or POPC/PG (20 mol%)/PS (4 mol%) (negatively charged membranes, open symbols). Acceptor LUVs A were composed of POPC (squares), POPC/PG (5 mol%) (solid circles), or POPC/PG (20 mol%) (open circles). PS transport was mediated by wild-type Osh6 and the Osh6^Δ35^, Osh6^Δ42^, and Osh6^lidless^ (black, orange, blue, and green symbols, respectively). c_PS in LUVs D_= 2 μM (1 μM in the outer leaflet accessible for Osh6), *c*_total lipids in LUVs D_ = 50 μM, c_total lipids in LUVs A_ = 200 μM, c_Osh6_ = 250 nM, c_C2Lact-CFP_ = 50 nM. **D)** Visualization of PS transport from LUVs to GUVs. Donor LUVs were composed of either POPC/PS (5 mol%) or POPC/PG (15 mol%)/PS (5 mol%), whereas GUVs were composed of either POPC or POPC/PG (20 mol%). PS transferred to GUVs was detected by recruitment of the fluorescent PS biosensor C2_Lact_-CFP. GUVs contained DiD for visualization in the red fluorescence channel. c_total lipids in GUVs_ = 12,5 μM, c_total lipids in LUVs_ = 150 μM, c_PS in LUVs_= 7,5 μM, c_Osh6_ = 250 nM, c_C2Lact-CFP_ = 50 nM.

We next asked how these differences in PS removal affect transport between membranes (Fig. **2A,C**). These measurements were performed in the extraction regime, but donor LUVs contained 4 mol% PS, corresponding to a fourfold molar excess over Osh6. Under these conditions, depletion exceeding the binding capacity of Osh6 requires repeated cargo extraction and release and therefore reports cycling PS transport rather than a single extraction event.

For wild-type Osh6, PS transport from neutral donor membranes occurred readily toward neutral acceptors and was slightly enhanced by 5 mol% PG in the acceptor membrane (Fig. **2C**, solid symbols). In contrast, transport from negatively charged donor membranes to neutral acceptors was undetectable but was restored when the acceptor membrane was also negatively charged (Fig. **2C**, open symbols).

Osh6^Δ35^ and Osh6^Δ42^ showed a similar dependence on membrane charge. Notably, Osh6^Δ42^ efficiently transported PS between two negatively charged membranes (Fig. **2C**, open circles and upward-pointing triangles), despite showing no detectable PS removal from negatively charged donor membranes in the absence of an acceptor (Fig. **2B**). Thus, as observed for PI4P, the presence of a second membrane enables efficient lipid transfer under conditions in which removal from the donor membrane alone is inefficient. In contrast, Osh6^lidless^ behaved differently: PS transport was markedly reduced and was particularly inefficient when negatively charged membranes were involved.

Together, these results reveal a common feature of PS and PI4P transport: lipid embedded in a membrane from which Osh6 alone removes little cargo can nevertheless undergo substantial intermembrane transfer when a suitable acceptor membrane is present. This suggests that the opposing membrane can facilitate cargo removal from the donor membrane and raises the possibility that membrane tethering contributes directly to this process.

To visualize the membrane-charge dependence of PS transfer, donor LUVs containing either POPC/PS (5 mol%) or POPC/PG (15 mol%)/PS (5 mol%) were mixed with neutral POPC or negatively charged POPC/PG (20 mol%) GUVs (Fig. **2D**). After addition of wild-type Osh6, transferred PS was visualized by recruitment of C2_Lact_-CFP to the GUV membrane. Consistent with the FCCS measurements, PS transfer from negatively charged donor membranes was readily detected only when the acceptor GUVs were also negatively charged. Because C2_Lact_-CFP binding depends strongly on membrane composition, this assay was used only for qualitative visualization of PS transfer.

### Membrane tethering correlates with acceptor-dependent lipid removal

The results above show that efficient lipid transfer can occur even under conditions in which little lipid removal is detected from an isolated donor membrane. We therefore asked whether the second membrane facilitates lipid removal by allowing Osh6 to engage donor and acceptor membranes simultaneously. Osh6 has previously been shown to tether membranes (11) and, in cells, interacts with the ER membrane tether Ist2. We therefore examined whether Osh6 membrane tethering correlates with acceptor-dependent lipid removal and how it is affected by progressive shortening of the N-terminal lid.

Membrane tethering was measured by FCCS using two populations of differently labeled LUVs (Fig. **3A**). Tethering results in co-diffusion of vesicles through the detection volume and therefore an increase in *G*_cc_(0). This assay detects membrane contacts that persist sufficiently long to produce measurable co-diffusion but does not require stable vesicle aggregation.

**Figure 3.**
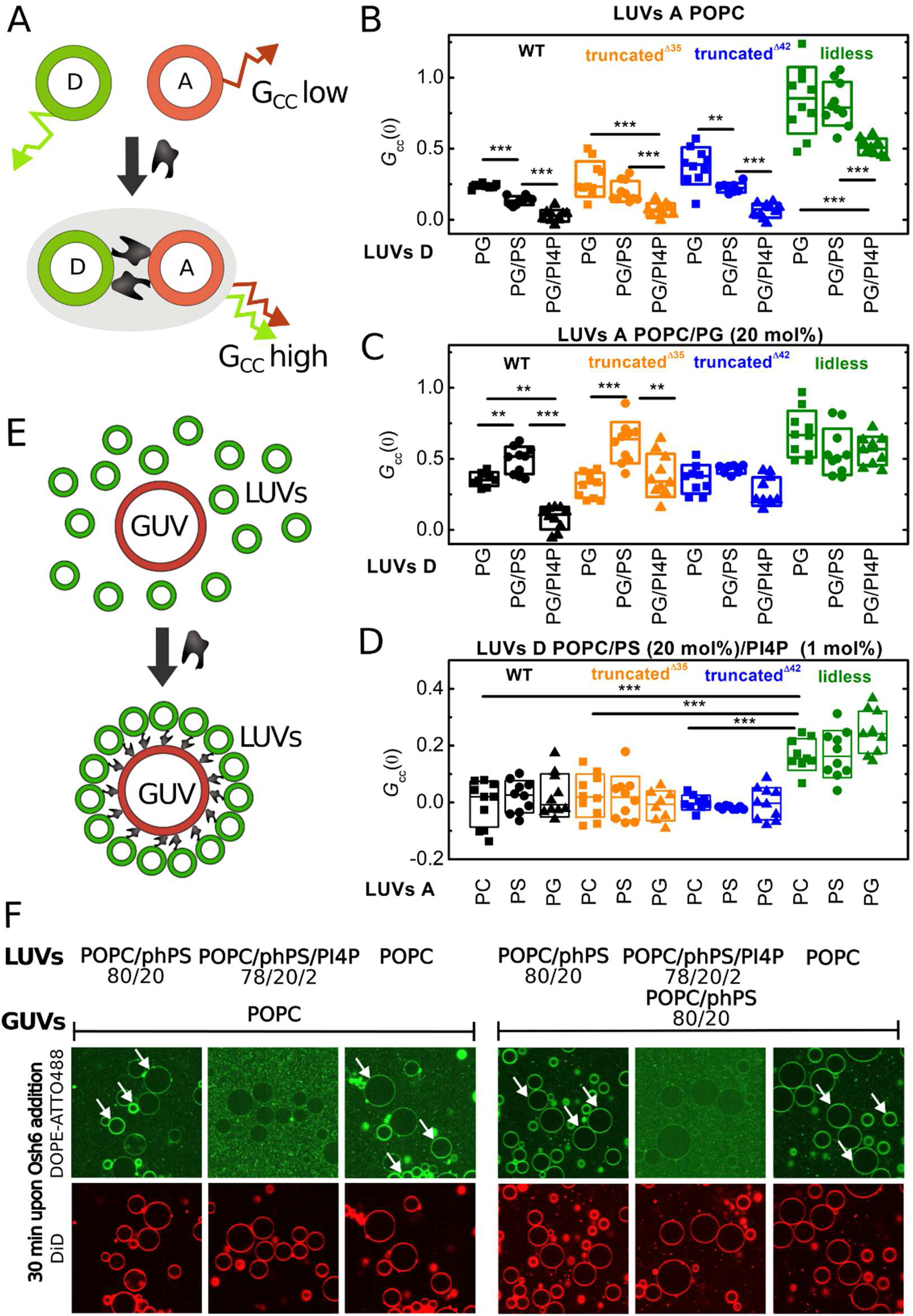
Membrane tethering mediated by wild-type Osh6 and N-terminal truncation mutants. **A)** Scheme of the FCCS assay used to monitor membrane tethering between donor LUVs (LUVs D, green) and acceptor LUVs (LUVs A, red). Independent diffusion of the two vesicle populations produces little cross-correlation, whereas Osh6-mediated tethering results in co-diffusion and an increased zero-time cross-correlation amplitude, *G*_cc_(0). **B–C)** Membrane tethering quantified by *G*_cc_(0). LUVs A were composed of either neutral POPC (**B**) or negatively charged POPC/PG (20 mol%) (**C**). LUVs D were composed of POPC/PG (20 mol%; squares), POPC/PG (15 mol%)/PS (5 mol%; circles), or POPC/PG (15 mol%)/PI4P (5 mol%; upward-pointing triangles). Wild-type Osh6 and the Osh6^Δ35^, Osh6^Δ42^, and Osh6^lidless^ mutants are shown in black, orange, blue, and green, respectively. Total lipid concentrations were 50 μM in both LUVs D and LUVs A; Osh6 concentration was 250 nM. **D)** Membrane tethering under conditions corresponding to the PI4P removal experiments in Fig. 1D, quantified by *G*_cc_(0). LUVs D were composed of POPC/PS (15 mol%)/PI4P (5 mol%), and LUVs A were composed of POPC (squares), POPC/PS (20 mol%; circles), or POPC/PG (20 mol%; upward-pointing triangles). Wild-type Osh6 and the Osh6^Δ35^, Osh6^Δ42^, and Osh6^lidless^ mutants are shown in black, orange, blue, and green, respectively. Total lipid concentrations were 50 μM in both LUVs D and LUVs A; Osh6 concentration was 250 nM. **E)** Scheme of the LUV–GUV tethering assay. Osh6-mediated tethering recruits DOPE-ATTO488-labeled LUVs (green) to DiD-labeled GUVs (red). **F)** Visualization of membrane tethering mediated by wild-type Osh6. DOPE-ATTO488-labeled LUVs of the indicated lipid compositions were incubated with DiD-labeled GUVs for 30 min in the presence of Osh6. LUVs contained either POPC, POPC/PS (20 mol%), POPC/PS (15 mol%)/PI4P (5 mol%), or POPC/1,2-diphytanoyl-sn-glycero-3-phosphoserine (phPS) (20 mol%). White arrows indicate GUVs decorated with tethered LUVs. Total lipid concentrations were 100 μM in LUVs and 12.5 μM in GUVs; Osh6 concentration was 100 nM.

We first measured tethering between neutral POPC LUVs A and negatively charged LUVs D containing POPC/PG (20 mol%), with 5 mol% PG replaced by either PS or PI4P (Fig. **3B**). With wild-type Osh6, Osh6^Δ35^, and Osh6^Δ42^, tethering was readily detected with cargo-free PG-containing LUVs D. Incorporation of either PS or PI4P reduced tethering, with PI4P producing the stronger effect. In contrast, Osh6^lidless^ displayed the highest tethering signal, consistent with its enhanced membrane association and impaired dissociation.

We next examined tethering when both membranes were negatively charged (Fig. **3C**). LUVs A contained POPC/PG (20 mol%), while the same set of negatively charged LUVs D as in Fig. **3B** was used. For wild-type Osh6 and Osh6^Δ35^, PS now enhanced rather than reduced tethering, whereas PI4P suppressed tethering under both membrane conditions. Tethering by Osh6^Δ42^ and Osh6^lidless^ was largely independent of donor cargo composition.

Thus, PS reduced tethering when only one membrane was negatively charged but enhanced tethering when both membranes were negatively charged. Notably, the latter conditions also support efficient PS transport despite inefficient PS removal from the donor membrane alone (Fig. **2**). The correspondence between enhanced tethering and efficient transport under conditions unfavorable for donor-alone PS removal is consistent with membrane tethering facilitating PS transfer. Tethering under similar membrane conditions was independently examined using the LUV–GUV system (Fig. **S3**). Wild-type Osh6 promoted recruitment of fluorescently labeled LUVs to the GUV surface, and the dependence of this recruitment on membrane charge and PS was consistent with the tethering behavior observed by FCCS. Thus, two independent experimental approaches support the ability of Osh6 to bridge membranes under conditions relevant to PS transport.

We next examined tethering under membrane conditions corresponding to the acceptor-dependent PI4P removal experiment in Fig. **1D**. LUVs D contained both PS and PI4P, while LUVs A had the same compositions as those used in the PI4P removal assay (Fig. **3D**). Under these conditions, substantial tethering was detected only for Osh6^lidless^, whereas wild-type Osh6 and the shorter truncation mutants showed little detectable tethering. Thus, Osh6^lidless^ retains pronounced tethering under the same membrane conditions in which its PI4P removal becomes strongly dependent on the presence of an acceptor membrane.

We further used the LUV–GUV system to examine how cargo identity affects the membrane-bridged state (Fig. **3E,F**). Recruitment of DOPE-ATTO488-labeled LUVs to the surface of DiD-labeled GUVs indicated tethering between the two membrane populations. Tethering was readily observed between neutral membranes, membranes of mixed charge, and negatively charged membranes containing 1,2-diphytanoyl-sn-glycero-3-phosphoserine (phPS), a PS analogue that is recognized by Osh6 but is not extracted (28). phPS was used to obtain a persistent tethered state in the presence of a recognized but non-transferable cargo, as PS-containing membranes did not maintain detectable tethering sufficiently long to test the effect of PI4P. Notably, incorporation of PI4P into the phPS-containing membrane abolished detectable tethering.

The contrasting effects of non-transferable phPS and PI4P suggest that productive PI4P extraction destabilizes the tethered state. Rather than representing a stable end state, membrane tethering may therefore constitute a transient intermediate during lipid transfer. Together with the FCCS measurements, these observations support a “kiss-and-ride” mechanism in which Osh6 transiently bridges two membranes during lipid transfer, while productive PI4P extraction destabilizes the bridged state and promotes Osh6 dissociation.

### Kinetic dissection of PI4P and PS transport

The results above suggest that the presence of an acceptor membrane can enable efficient lipid transfer under conditions in which little cargo removal is detected from the donor membrane alone. Moreover, PI4P strongly suppresses detectable membrane tethering, whereas non-transferable phPS does not (Fig. **3**). Together, these observations support a “kiss-and-ride” mechanism in which Osh6 transiently engages both membranes, facilitating PI4P extraction within a membrane-bridged state that is destabilized upon productive cargo extraction. Because such transient intermediates are difficult to visualize directly, we sought kinetic evidence for their contribution to lipid transport.

We first compared the rate of PI4P transport with the rate of PI4P release from preloaded Osh6 under matched membrane conditions (Fig. **4A**). Donor membranes (LUVs D) were either neutral or negatively charged through incorporation of PS or PG, whereas acceptor membranes (LUVs A) covered a range of neutral and negatively charged compositions. For transport measurements, LUVs D additionally contained 2 mol% PI4P.

**Figure 4.**
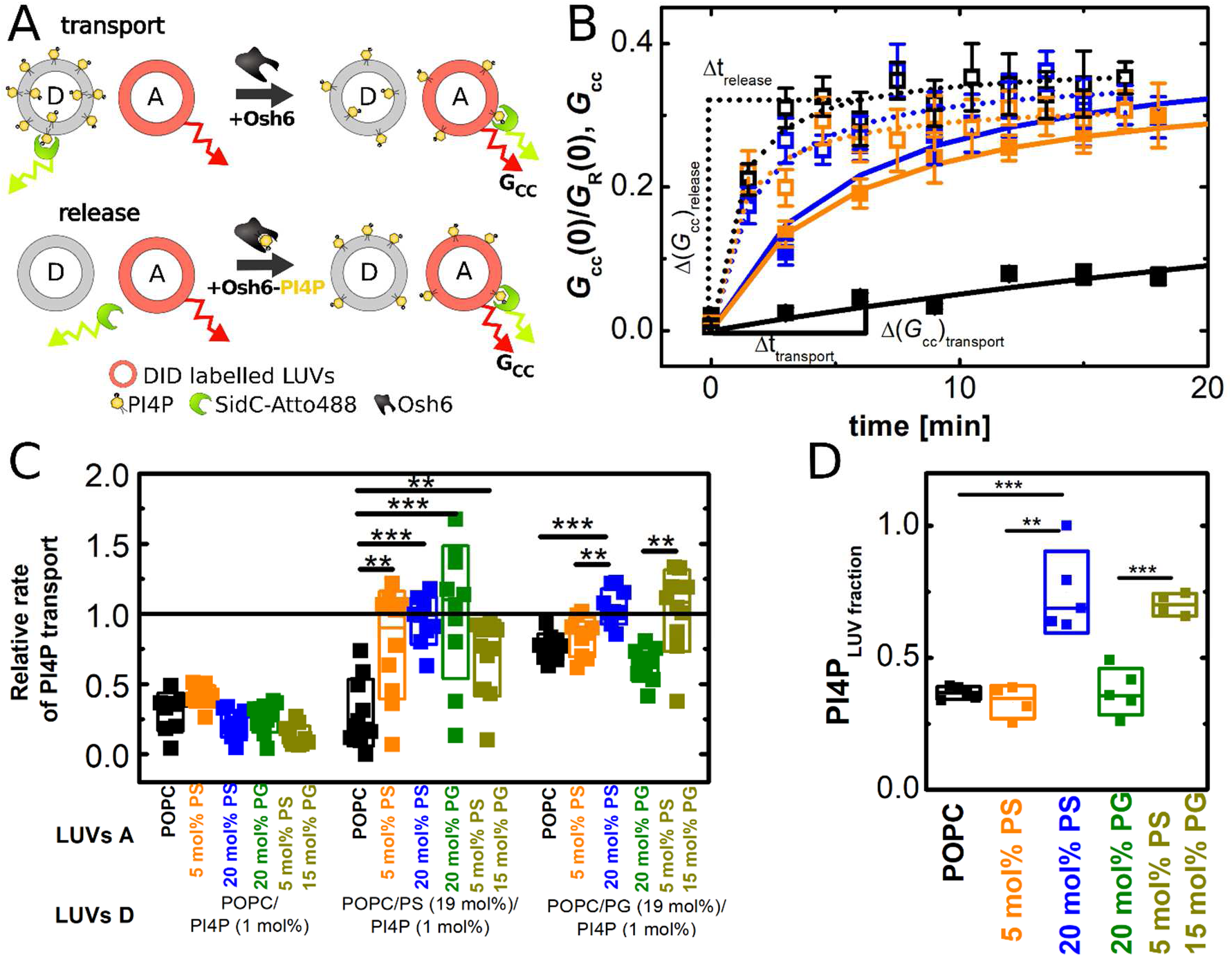
Kinetic comparison of PI4P transport and release from Osh6. **A)** Schematic representation of the transport–release experiment. In the transport experiment, PI4P transfer from donor LUVs (LUVs D) to acceptor LUVs (LUVs A) was monitored as a time-dependent increase in the FCCS signal, *G*_cc_, between the PI4P biosensor SidC-Atto488 and DiD in acceptor LUVs. In the release experiment, PI4P-loaded Osh6 was added to an identical mixture of PI4P-free donor and acceptor LUVs. The amount of PI4P preloaded into Osh6 was equivalent to the total transportable PI4P in the transport experiment. Transport and release were measured under identical membrane conditions. **B)** Representative PI4P transport (solid symbols) and release (open symbols) kinetics, expressed as the time-dependent increase in *G*_cc(_0). LUVs D were composed of POPC/PI4P (2 mol%; black), POPC/PG (15 mol%)/PI4P (2 mol%; orange), or POPC/PS (15 mol%)/PI4P (2 mol%; blue); PI4P was omitted from LUVs D in the corresponding release experiments. LUVs A were composed of POPC/PS (20 mol%). Transport and release rates were determined from the change in *G*_cc_(0) over the indicated time interval. **C)** Relative PI4P transport rates, calculated as the ratio of the transport rate to the release rate, for different donor (LUVs D) and acceptor (LUVs A) membrane compositions. Donor LUVs were composed of POPC/PI4P (2 mol%) (left), POPC/PS (15 mol%)/PI4P (2 mol%) (middle), and POPC/PG (15 mol%)/PI4P (2 mol%) (right). Acceptor LUVs were composed of POPC (black), POPC/PS (5 mol%) (orange), POPC/PS (20 mol%) (blue), POPC/PG (20 mol%) (green), and POPC/PS (5 mol%)/PG (15 mol%) (dark yellow). c_total lipids in LUVs D_ = 50 μM, c_total lipids in LUVs A_ = 50 μM, c_PI4P in LUVs D_ = 1 μM, c_Osh6_ = 250 nM, c_SidC-Atto488_ = 80 nM. **D)** Equilibrium partitioning of PI4P between Osh6 and LUVs of defined composition, expressed as the fraction of PI4P associated with LUVs. LUVs were composed of POPC, POPC/PS (5 mol%), POPC/PS (20 mol%), or POPC/PS (5 mol%)/PG (15 mol%). Total lipid concentration was 50 μM; PI4P-loaded Osh6 and SidC-Atto488 concentrations were 125 nM and 80 nM, respectively.

PI4P transport was monitored by FCCS in the transport regime, with labeled LUVs A and unlabeled LUVs D. The increase in *G*_cc_(0), reflecting PI4P accumulation in LUVs A, was followed over time to determine the transport rate (Fig. **4B,C**). These kinetics were compared with PI4P release from preloaded Osh6 into the same donor–acceptor membrane mixtures, except that LUVs D contained no PI4P. Thus, the release assay measured PI4P release from Osh6 and accumulation in the acceptor membrane without the preceding cargo-acquisition step.

To compare these processes, we defined the relative transport rate as the ratio of the transport rate to the PI4P release rate measured under matched membrane conditions (Fig. **4B**):

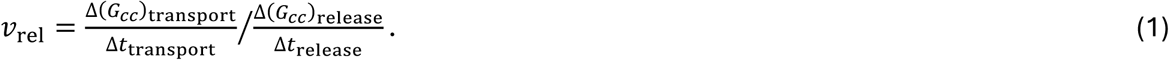

For neutral donor membranes, the relative transport rate remained below 0.5 with all acceptor compositions tested (Fig. **4C**). In contrast, with negatively charged donor membranes containing either PS or PG, the transport rate approached the PI4P release rate when the acceptor membrane was also negatively charged. Most notably, this behavior was observed with PS-rich donor membranes, from which little PI4P removal was detected in the absence of an acceptor (Fig. **1D**). Thus, PI4P acquisition from an isolated PS-rich donor membrane is strongly disfavored, yet in the presence of a suitable opposing membrane overall transport proceeds at a rate approaching PI4P release from preloaded Osh6.

Importantly, this effect could not be explained simply by the cargo-loaded state of Osh6. No detectable PI4P removal from PS-rich donor membranes was observed even when Osh6 was preloaded with PS (Fig. **S1**). Thus, PS loading alone cannot account for efficient PI4P acquisition under these conditions. Instead, the presence of the opposing membrane changes the conditions for PI4P removal from the donor.

To assess the contribution of membrane-dependent PI4P partitioning, we determined the equilibrium distribution of PI4P between Osh6 and PI4P-free LUVs of defined composition (Fig. **4D**). PI4P-loaded Osh6 was added to the LUVs, and the fraction of PI4P transferred to the membranes at equilibrium was quantified. PI4P showed the strongest equilibrium preference for negatively charged, PS-containing membranes relative to Osh6. However, because transport and release were compared under matched membrane conditions, differences in PI4P partitioning alone cannot account for the observed relationship between the two rates.

We next asked whether PS shows a similar kinetic behavior. As shown in Figs. **2** and **3**, little PS removal was detected from negatively charged donor membranes in the absence of an acceptor, whereas efficient PS transport occurred between negatively charged membranes and was associated with enhanced membrane tethering.

Using the same experimental strategy as for PI4P, we compared PS transport with PS release from preloaded Osh6 under matched membrane conditions (Fig. **5A**). Donor membranes were either neutral POPC or negatively charged POPC/PG (19 mol%), each containing 1 mol% PS. Acceptor membranes were neutral or negatively charged and either contained PI4P or lacked it.

**Figure 5.**
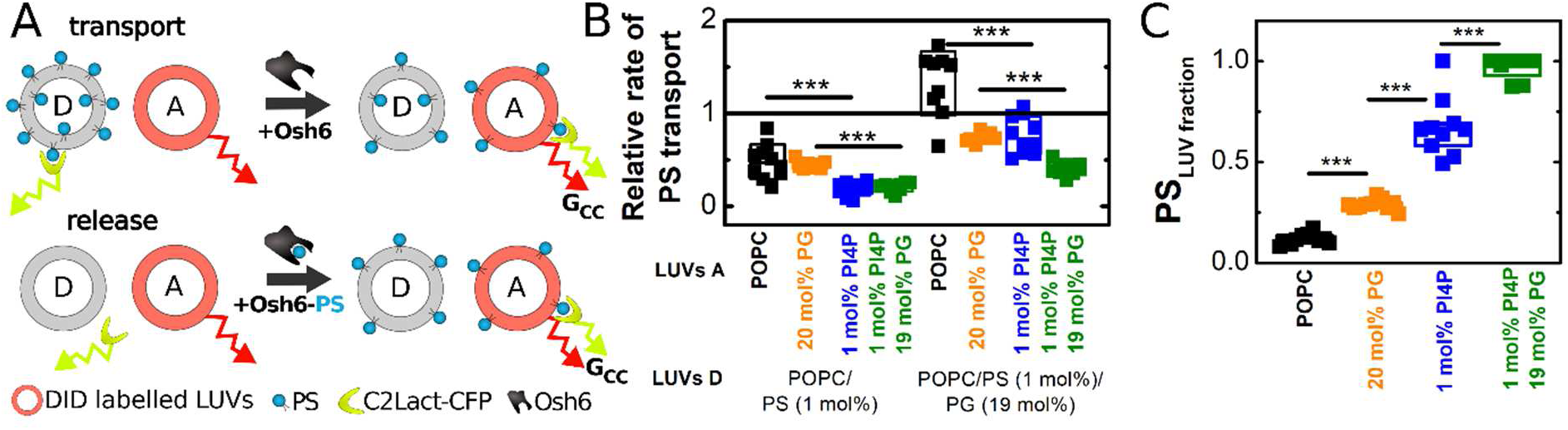
Kinetic comparison of PS transport and release from Osh6. **A)** Schematic representation of the PS transport–release experiment. In the transport experiment, PS transfer from donor LUVs (LUVs D) to acceptor LUVs (LUVs A) was monitored as a time-dependent increase in the FCCS cross-correlation amplitude, *G*_cc_(0), between the PS biosensor C2_Lact_-CFP and DiD in LUVs A. In the release experiment, PS-loaded Osh6 was added to the same donor–acceptor membrane mixture, except that PS was omitted from LUVs D. Transport and release were thus measured under matched membrane conditions. **B)** Relative PS transport rates, calculated as the ratio of the transport rate to the release rate under matched membrane conditions, for different combinations of LUVs D and LUVs A. LUVs D were composed of POPC/PS (1 mol%) or POPC/PG (19 mol%)/PS (1 mol%). LUVs A were composed of POPC (black), POPC/PG (20 mol%; orange), POPC/PI4P (1 mol%; blue), or POPC/PG (19 mol%)/PI4P (1 mol%; green). Total lipid concentrations were 50 μM in both LUVs D and LUVs A. PS concentration in LUVs D was 500 nM; Osh6 and C2_Lact_-CFP concentrations were 250 nM and 50 nM, respectively. **C)** Equilibrium partitioning of PS between Osh6 and LUVs of defined composition, expressed as the fraction of PS associated with LUVs. LUVs were composed of POPC, POPC/PG (20 mol%), POPC/PI4P (1 mol%), or POPC/PG (19 mol%)/PI4P (1 mol%). Total lipid concentration was 50 μM; PS-loaded Osh6 and C2_Lact-_CFP concentrations were 125 nM and 50 nM, respectively.

Transport from negatively charged donor membranes approached the PS release rate, whereas transport from neutral donor membranes remained substantially slower (Fig. **5B**). Thus, as for PI4P, PS removal from an isolated negatively charged donor membrane is inefficient, yet the presence of an appropriate opposing membrane allows transport to proceed at a rate approaching cargo release from preloaded Osh6. Together with the enhanced tethering observed under these membrane conditions (Fig. **3C**), this kinetic behavior is consistent with engagement of the opposing membrane facilitating PS extraction.

We also determined the equilibrium distribution of PS between Osh6 and LUVs of defined composition (Fig. **5C**). PS partitioning depended strongly on membrane composition: neutral POPC membranes accepted substantially less PS than negatively charged membranes, with the strongest PS partitioning observed toward PI4P-containing membranes.

Thus, both Osh6 cargoes show a similar relationship between donor-alone removal and transport kinetics: membrane conditions that strongly disfavor cargo removal from an isolated donor can nevertheless support rapid transport in the presence of a suitable opposing membrane. Their effects on the tethered state, however, differ. PI4P suppresses detectable membrane tethering, consistent with productive PI4P extraction destabilizing a transient bridged state, whereas pronounced tethering persists under conditions supporting efficient PS transport between negatively charged membranes. The lifetime of the membrane-bridged state therefore appears to depend on both cargo identity and membrane composition.

### Complementation of yeast phenotypes by Oshc mutants

Based on our *in vitro* results, we expected the progressive N-terminal truncation of Osh6 to differentially affect Osh-dependent cellular functions. Donor-alone removal of both PI4P and PS was strongly impaired by N-terminal truncation, yet PI4P transport remained largely preserved even in Osh6^lidless^, whereas PS transport was strongly reduced. We therefore asked whether these distinct *in vitro* phenotypes would also be reflected *in vivo* by testing the ability of the Osh6 mutants to complement two Osh-dependent yeast phenotypes.

We first examined a phenotype dependent on physiologically relevant PS transport using the *dpl1Δ osh6Δ osh7Δ psd1Δ* strain (35, 36). In this strain, conversion of PS to phosphatidylethanolamine (PE) by Psd2 depends on PS transport mediated by Osh6 and Osh7 (18, 35, 37, 38). Consequently, loss of Osh6 and Osh7 strongly impairs growth in the absence of ethanolamine, providing a functional readout of PS transport.

Expression of wild-type Osh6 restored growth in the absence of ethanolamine (Fig. **6A**). Osh6^Δ35^ complemented the growth defect similarly to wild-type Osh6, whereas Osh6^Δ42^ showed intermediate complementation. In contrast, Osh6^lidless^ failed to restore ethanolamine prototrophy, consistent with severe impairment of physiologically relevant PS transport upon complete removal of the lid.

**Figure 6.**
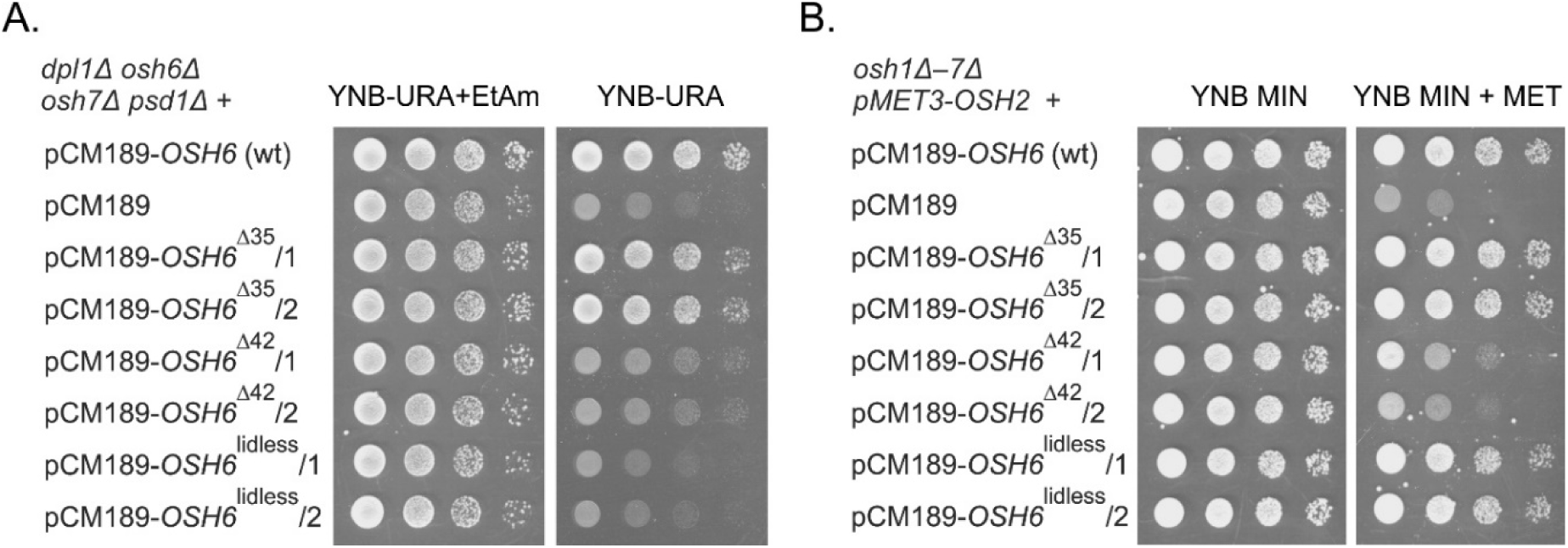
Functional complementation by wild-type Osh6 and N-terminal truncation mutants in yeast. **A)** Complementation of ethanolamine-dependent growth of the *dpl1Δ osh6Δ osh7Δ psd1Δ* strain by Osh6 N-terminal truncation mutants. Wild-type *OSH6* and the indicated truncation mutants (Osh6^Δ35^, Osh6^Δ42^, and Osh6^lidless^) were expressed from the yeast expression vector pCM189. Cells carrying the empty pCM189 vector served as a negative control. Cultures were pre-grown in YNB-URA medium supplemented with ethanolamine (EtAm), serially diluted 10-fold, spotted onto YNB-URA plates with or without EtAm, and incubated at 30 °C for 2 days. Two independent transformants were analyzed for each construct. **B)** Complementation of the growth defect of the *osh1Δ–7Δ pMET3-OSH2* strain by Osh6 N-terminal truncation mutants. Cells were transformed with pCM189 plasmids expressing wild-type *OSH6* or the indicated truncation mutants under the control of the tetO-CYC1 promoter. Cells carrying the empty pCM189 vector served as a negative control, and cells expressing wild-type *OSH6* served as a positive control. Cultures were pre-grown in YNB-MIN medium, serially diluted 10-fold, spotted onto YNB-MIN plates with or without methionine (100 mg/L), and incubated at 30 °C for 2 days. Two independent transformants were analyzed for each construct.

We next asked whether the Osh6 mutants could substitute for the essential cellular functions of the entire Osh protein family. Although none of the seven yeast Osh proteins is individually essential, simultaneous deletion of all *OSH* genes is lethal (39). We therefore used an *osh1Δ–7Δ pMET3-OSH2* strain in which viability is maintained by expression of *OSH2* from the methionine-repressible *MET3* promoter (39). Loss of Osh function is associated with severe disruption of phosphoinositide homeostasis, including accumulation of PI4P at the plasma membrane (40, 41). Complementation in this system therefore provides a stringent functional test of the ability of Osh6 mutants to substitute for endogenous Osh proteins *in vivo*. Upon addition of methionine, *OSH2* expression is repressed and growth becomes dependent on the introduced Osh6 construct (Fig. **6B**).

Osh6^Δ35^ and Osh6^lidless^ restored growth under these conditions to levels comparable to wild-type Osh6. Thus, despite complete removal of the N-terminal lid and its strong effect on PS transport, Osh6^lidless^ retains the ability to substitute for essential cellular functions of the Osh protein family. In contrast, Osh6^Δ42^ supported only limited growth.

Together, these two cellular assays reveal distinct functional consequences of N-terminal lid shortening. Progressive truncation impaired the PS-dependent phenotype, culminating in loss of complementation by Osh6^lidless^, whereas the ability to support growth in the absence of endogenous Osh proteins did not follow the same trend: Osh6^lidless^ retained substantial activity despite the weaker complementation by Osh6^Δ42^. Thus, complete lid removal does not simply render Osh6 non-functional *in vivo*, but differentially affects distinct Osh-dependent cellular functions.

## Materials and Methods

### Protein expression and purification

Genes encoding Osh6, its N-terminal truncation mutants, SidC, and C2_Lact_-CFP were cloned into a modified pHIS2 expression vector containing an N-terminal His6 tag followed by a tobacco etch virus (TEV) protease cleavage site. To enable site-specific fluorescent labeling, a cysteine-free Osh6 construct (C62S/C162S/C389S) and a single-cysteine variant (I241C) were generated by site-directed mutagenesis. Recombinant proteins were produced in *Escherichia coli* BL21 Star cells following previously established procedures (42, 43).

Cells were harvested by centrifugation, resuspended in lysis buffer (50 mM Tris, pH 8.0, 300 mM NaCl, 20 mM imidazole, 3 mM β-mercaptoethanol, and 10% glycerol), and disrupted by sonication. Cell debris was removed by centrifugation, and the clarified lysate was applied to Ni²⁺ affinity resin (Macherey–Nagel). Bound proteins were eluted with the same buffer containing 300 mM imidazole, followed by TEV protease cleavage to remove the His6 tag.

Proteins were further purified by size-exclusion chromatography on a HiLoad 16/600 Superdex 75 pg column (Cytiva) equilibrated with 20 mM Tris (pH 7.4), 300 mM NaCl, 3 mM β-mercaptoethanol, and 10% glycerol. Osh6 constructs underwent an additional purification step by ion-exchange chromatography on a HiTrap SP HP column (Cytiva).

For preparation of lipid-loaded Osh6, the protein was mixed with LUVs composed of POPC/PS/DiD (79.7:20:0.3 mol/mol/mol) or POPC/PI4P/DiD (89.7:10:0.3 mol/mol/mol), prepared at a total lipid concentration of 10 mM. The volume of LUVs added was adjusted such that PS or PI4P was present at a threefold molar excess over Osh6. The mixtures were incubated overnight at 4 °C. DiD was included in the LUVs to facilitate identification of the vesicle-containing fractions during size-exclusion chromatography. Lipid-loaded Osh6 was subsequently separated from the LUVs on a HiLoad Superdex 200 pg column (Cytiva) equilibrated with 20 mM Tris (pH 7.4), 300 mM NaCl, 3 mM β-mercaptoethanol, and 10% glycerol.

For fluorescent labeling, SidC and the single-cysteine Osh6 variant were buffer-exchanged into PBS and incubated overnight at 4 °C with a threefold molar excess of Atto488-maleimide. Unreacted fluorophore was removed by size-exclusion chromatography on a Superdex 200 pg column (Cytiva) using 20 mM Tris (pH 7.4), 300 mM NaCl, 3 mM β-mercaptoethanol, and 10% glycerol as the running buffer.

### Lipids and other chemicals

All lipids, including brain PI4P, were obtained from Avanti Polar Lipids (Alabaster, AL, USA) and used without further purification. Atto488-labeled DOPE was obtained from ATTO-TEC (Siegen, Germany), whereas DiD and all other chemicals were purchased from Sigma-Aldrich (St. Louis, MO, USA).

### LUV formation

Lipids dissolved in organic solvents were mixed at the desired molar ratios, dried under a stream of nitrogen, and further desiccated under vacuum for at least 2 h to remove residual solvent. Lipid films were hydrated to a final lipid concentration of 1 mM in LUV buffer (40 mM imidazole, pH 7.4, 150 mM NaCl, 3 mM β-mercaptoethanol, and 1 mM EDTA). For GUV experiments, lipids were hydrated in GUV buffer (25 mM Tris, pH 8.0, 10 mM MgCl₂, 20 mM imidazole, 261.5 mM NaCl, and 2 mM β-mercaptoethanol). Lipid suspensions were vortexed until fully resuspended, subjected to one freeze–thaw cycle, and extruded 41 times through 50-nm polycarbonate filters using a mini-extruder (Avestin, Ottawa, Canada). LUVs were used immediately or stored under nitrogen at room temperature for up to 24 h (44).

### GUV preparation

GUVs were prepared by electroformation. Lipids dissolved in organic solvents were mixed at the desired molar ratios together with DiD (1:6500 mol/mol relative to total lipid). A total of 100 nmol of lipids in 100 μL chloroform was spread evenly onto two recessed titanium electrodes and dried under vacuum for at least 1 h to remove residual solvent. The lipid-coated electrodes were assembled using a Parafilm spacer to form the electroformation chamber, which was filled with 1 mL of 600 mM sucrose adjusted to match the osmolality of the GUV buffer and sealed with Parafilm. Electroformation was performed at 47 °C using the following protocol: 10 Hz at 1.0 V for 15 min, followed by 10 Hz at 1.5 V for 90 min and finally 2 Hz at 1.5 V for 20 min. GUVs were collected immediately after electroformation and used for microscopy.

### Microscopy

FCCS measurements were performed on a Leica SP8 confocal microscope (Leica, Mannheim, Germany) equipped with a 63× water-immersion objective (N.A. 1.2), synchronized pulsed laser sources, and hybrid HyD detectors. DiD was excited using the 640-nm line of the white-light laser (Coherent, Santa Clara, CA, USA), whereas CFP and Atto488 were excited using 440-nm and 470-nm pulsed diode lasers (PicoQuant, Berlin, Germany), respectively. The 440/640-nm and 470/640-nm laser pairs were operated in pulsed interleaved excitation (PIE) mode at overall repetition rates of 40 and 20 MHz, respectively. PIE enabled fluorescence photons originating from different excitation lasers to be distinguished based on both their emission spectra and arrival times, thereby minimizing spectral cross-talk between detection channels. Correlation analysis was performed using custom MATLAB scripts (MathWorks, Natick, MA, USA).

Confocal imaging was performed on a Zeiss LSM 980 microscope (Carl Zeiss, Oberkochen, Germany) equipped with a 40× water-immersion objective (N.A. 1.2). DiD, Atto488, and CFP were excited using 639 nm, 488 nm and 445 nm laser lines, respectively. Image analysis was performed using FIJI/ImageJ.

### FCCS assay

The FCCS assay was used to monitor lipid removal, lipid transport, membrane tethering, and lipid release under the experimental conditions summarized in Table **1**. Donor LUVs (LUVs D) and, where applicable, acceptor LUVs (LUVs A) were mixed in the proportions specified in Table **1** and transferred to BSA-coated four-chamber 35-mm glass-bottom dishes (Cellvis, USA) containing LUV buffer supplemented with the appropriate lipid biosensor (50 nM C2_Lact_-CFP for PS assays or 80 nM SidC-Atto488 for PI4P assays). The final assay volume was adjusted to 200 μL with LUV buffer. A 200-s baseline was recorded before addition of Osh6. Measurements were initiated by addition of Osh6 or cargo-loaded Osh6, as appropriate for the individual assay, and data acquisition continued for an additional 15– 20 min.

**Table 1.**
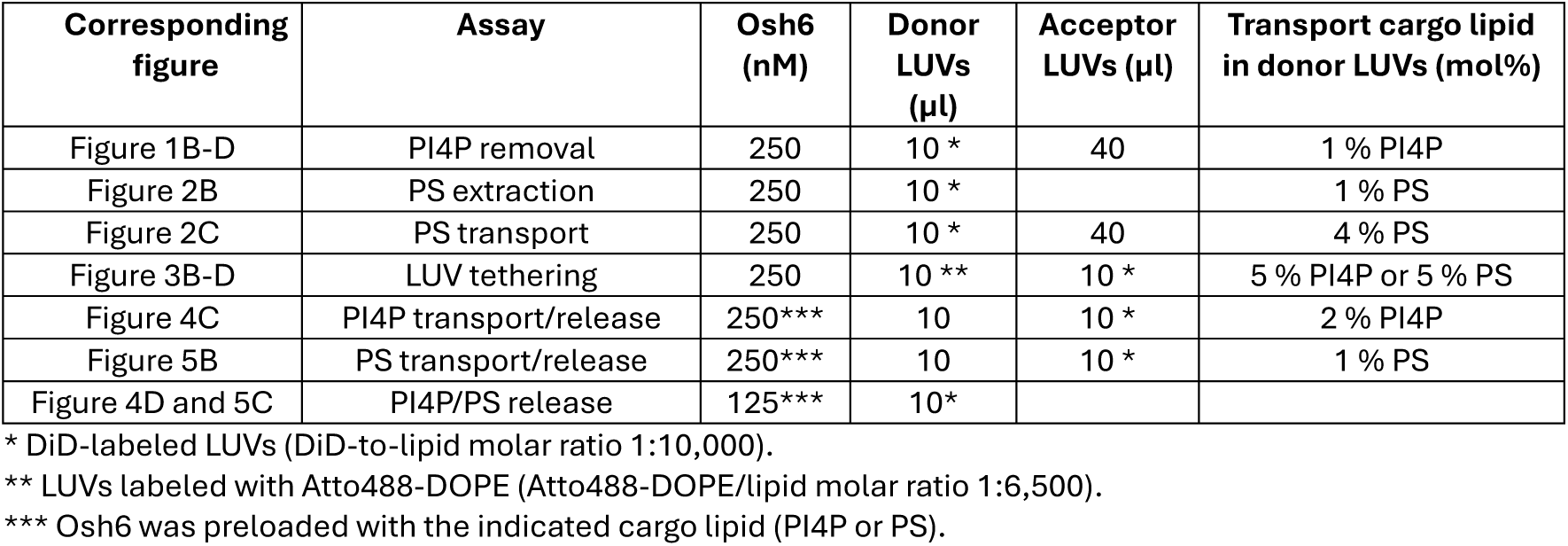
Experimental conditions used in individual LUV experiments and corresponding figures.

| Corresponding figure | Assay | Osh6 (nM) | Donor LUVs (μl) | Acceptor LUVs (μl) | Transport cargo lipid in donor LUVs (mol%) |
| --- | --- | --- | --- | --- | --- |
| Figure 1B-D | PI4P removal | 250 | 10 * | 40 | 1 % PI4P |
| Figure 2B | PS extraction | 250 | 10 * |  | 1 % PS |
| Figure 2C | PS transport | 250 | 10 * | 40 | 4 % PS |
| Figure 3B-D | LUV tethering | 250 | 10 ** | 10 * | 5 % PI4P or 5 % PS |
| Figure 4C | PI4P transport/release | 250*** | 10 | 10 * | 2 % PI4P |
| Figure 5B | PS transport/release | 250*** | 10 | 10 * | 1 % PS |
| Figure 4D and 5C | PI4P/PS release | 125*** | 10* |  |  |

\* DiD-labeled LUVs (DiD-to-lipid molar ratio 1:10,000).
\*\* LUVs labeled with Atto488-DOPE (Atto488-DOPE/lipid molar ratio 1:6,500).
\*\*\* Osh6 was preloaded with the indicated cargo lipid (PI4P or PS).

For lipid-removal assays, donor LUVs were labeled with DiD and the cargo lipid was detected using C2_Lact_-CFP for PS or SidC-Atto488 for PI4P. Binding of the biosensor to cargo-containing donor LUVs generated correlated fluorescence fluctuations, and removal of cargo from the donor membrane was detected as a decrease in the zero-lag cross-correlation amplitude, *G*_cc_(0).

For PS transport measurements shown in Fig. **2C**, transport was monitored through depletion of PS from donor LUVs. Donor LUVs contained 4 mol% PS, corresponding to 2 μM total PS and approximately 1 μM PS in the outer leaflet accessible to Osh6. Because accessible PS was present in fourfold molar excess over Osh6 (250 nM), depletion exceeding the cargo-binding capacity of Osh6 required repeated cycles of PS extraction and release and therefore reported sustained PS transport in the presence of acceptor membranes. Quantification was performed using membrane-composition-specific calibration curves as described below.

For membrane-tethering measurements, the two LUV populations were labeled separately with DiD and Atto488-DOPE. Osh6-mediated association of the two vesicle populations resulted in their co-diffusion through the confocal volume and was quantified by the resulting increase in *G*_cc_(0).

### Transport–release measurements

For the kinetic comparison of lipid transport and cargo release (Figs. **4** and **5**), transport and release were measured under matched membrane conditions. In the transport experiment, cargo-containing donor LUVs and cargo-free acceptor LUVs were mixed, and transport was initiated by addition of Osh6. Transfer of PI4P or PS to DiD-labeled acceptor LUVs was monitored as a time-dependent increase in *G*_cc_(0) between DiD and the corresponding fluorescent lipid biosensor.

In the corresponding release experiment, Osh6 preloaded with PI4P or PS was added to the same donor–acceptor membrane mixture, except that the cargo lipid was omitted from donor LUVs. The increase in *G*_cc_(0) therefore reflected release of preloaded cargo from Osh6 and its accumulation in acceptor LUVs without a preceding cargo-extraction step. Transport and release rates were determined from the initial change in *G*_cc_(0) over the same time interval. Relative transport rates were calculated as the ratio of the transport rate to the corresponding cargo-release rate measured under matched membrane conditions.

### Estimation of lipid removal and transport rates

For PI4P and PS removal assays (Fig. **1** and Fig. **2B**), the initial rate of cargo removal was calculated from the time-dependent decrease in the normalized FCCS readout according to Eq. **2**:

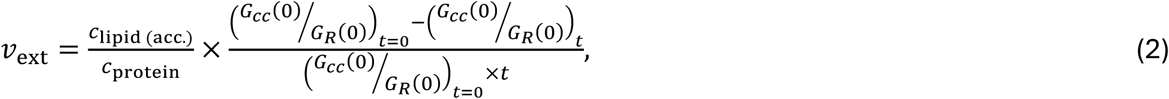

where *c*_lipid(acc.)_is the concentration of accessible cargo lipid in donor LUVs, *c*_protein_ is the concentration of Osh6, and (*G_cc_*(0)/*G_R_*(0))*_t_*_=0_ and (*G_cc_*(0)/*G_R_*(0))*_t_*represent the normalized FCCS parameter before addition of the transporter and at time *t*, respectively.

Accurate quantification of PS transport (Fig. **2C**), particularly between membranes differing in surface charge, required calibration of the PS biosensor C2_Lact_-CFP (34), whose response depends not only on the amount of PS but also on the membrane environment in which PS is detected. Calibration curves were therefore established separately for each membrane composition used in the study.

Calibration samples representing defined extents of PS transfer were prepared by gradually decreasing the PS content in donor LUVs while increasing the PS content in acceptor LUVs. FCCS measurements of these samples yielded calibration curves relating the normalized biosensor response to the fraction of transferred PS (Fig. **S2**). During transport experiments, the decrease in PS concentration in donor LUVs was converted to the corresponding extent of PS transfer using the appropriate calibration curve. Transport rates were then calculated according to Eq. **3**:

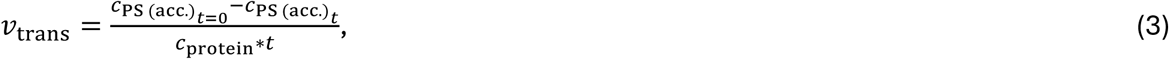

where *c*_PS(acc.),*t*=0_ and *c*_PS(acc.),*t*_ denote the concentrations of accessible PS before transporter addition and at time *t*, respectively.

### Determination of cargo partitioning

The equilibrium partitioning of PI4P and PS between Osh6 and membranes of defined composition (Figs. **4D** and **5C**) was determined in two sequential steps.

First, calibration curves were established for each membrane composition. DiD-labeled LUVs containing increasing amounts (0–0.5 mol%) of either PI4P or PS were prepared and mixed (10 μL) with the corresponding fluorescent biosensor (80 nM SidC-Atto488 for PI4P or 50 nM C2_Lact_-CFP for PS) in LUV buffer to a final volume of 200 μL. The normalized FCCS parameter was measured as a function of lipid concentration (Fig. **S4**), generating calibration curves relating the FCCS signal to the concentration of cargo lipid in each membrane environment.

In the second step, Osh6 preloaded with either PI4P or PS was mixed with cargo-free LUVs of the corresponding lipid composition. The increase in the FCCS signal was monitored until equilibrium was reached, and the plateau value was converted to the concentration of cargo lipid in the membrane using the appropriate calibration curve. From this value, the equilibrium distribution of cargo between the membrane and the Osh6 lipid-binding cavity was determined.

### GUV imaging

GUV-based assays were used to visualize Osh6-mediated lipid transport and membrane tethering under the experimental conditions summarized in Table **2**. GUVs and LUVs were mixed in the proportions indicated in Table **2** and transferred to BSA-coated glass-bottom chambers containing isosmotic GUV buffer supplemented with Osh6 and, for lipid-transfer assays, the appropriate fluorescent biosensor (80 nM SidC-Atto488 for PI4P assays or 50 nM C2_Lact_-CFP for PS assays). The final assay volume was adjusted to 200 μL with GUV buffer. Samples were incubated for the indicated times to allow Osh6-mediated lipid transfer or membrane tethering before confocal imaging. Control samples were prepared identically except that Osh6 was omitted.

**Table 2.** Experimental conditions used in individual GUV experiments and corresponding figures.

| Corresponding figure | Assay | Osh6 (nM) | GUV volume ($\mu\text{L}$) | LUV volume ($\mu\text{L}$) | Cargo lipid in donor LUVs (mol%) |
| --- | --- | --- | --- | --- | --- |
| Figure 1E | PI4P transport | 60 | 25* | 50 | 1 % PI4P |
| Figure 2D | PS transport | 250 | 25* | 30 | 5 % PS |
| Figure 3F | Tethering | 100 | 25* | 20** |  |

\*DiD-labeled GUVs (DiD-to-lipid molar ratio 1:5000).
\*\* LUVs labeled with Atto488 DOPE (Atto488 DOPE -to-lipid molar ratio 1:5000).

### Strains and culture conditions

The *Saccharomyces cerevisiae* strains used for functional complementation are listed below. Strain JRY6326 (**α** *leu2-3,112 ura3-52 his3Δ200 trp1-Δ901 suc2Δ9 osh1Δ::kanMX4 osh2Δ::kanMX4 osh3Δ::LYS2 osh4Δ::HIS3 osh5Δ::LEU2 osh6Δ::LEU2 osh7Δ::HIS3 TRP1*::pMET3*-OSH2*), used to assess complementation in the absence of all endogenous Osh proteins, was kindly provided by C. Beh (Simon Fraser University, Canada) (40). The *dpl1Δ osh6Δ osh7Δ psd1Δ* strain AWY722 (**α** *leu2 his3 lys2 ura3 dpl1Δ::LEU2 osh6Δ::kanMX osh7Δ::HisMX psd1Δ::natMX*) was obtained from the laboratory of C. Loewen (University of British Columbia, Canada) (35).

Unless otherwise indicated, yeast strains were cultured in yeast extract–peptone– dextrose (YEPD) medium containing 2% glucose. Media used for strains requiring ethanolamine were supplemented with 2 mM ethanolamine after sterilization. The *osh1Δ– 7Δ pMET3-OSH2* strain was maintained on YNB-MET medium (0.67% yeast nitrogen base, 2% glucose, containing all required amino acids and nucleobases with the exception of methionine) or on YNB-MIN medium (0.67% yeast nitrogen base without amino acids, 2% glucose) when required for plasmid maintenance. Strains carrying episomal plasmids were maintained and pre-cultured in synthetic minimal medium supplemented with the amino acids and nucleobases required for plasmid selection. Solid media contained 2% agar.

### Phenotypic testing

Functional complementation by Osh6 N-terminal truncation mutants was assessed by spot assays. Yeast cultures were grown overnight in selective medium appropriate for plasmid maintenance. Ten-fold serial dilutions were spotted onto agar plates, and growth was evaluated after 2 days of incubation at 30 °C.

For complementation of ethanolamine-dependent growth in the *dpl1Δ osh6Δ osh7Δ psd1Δ* strain, cultures were pre-grown in YNB-URA medium supplemented with ethanolamine and spotted onto YNB-URA plates with or without ethanolamine.

For complementation in the *osh1Δ–7Δ pMET3-OSH2* strain, cultures were pre-grown in YNB-MIN medium and spotted onto YNB-MIN plates with or without methionine (100 mg/L). Two independent transformants were analyzed for each construct.

### Plasmid constructio

Plasmid pCM189-*OSH6* (45)(kindly donated by M. Balážová, Centre of Biosciences, SAS, Slovakia) was generated by amplifying the *OSH6* coding sequence from yeast chromosomal DNA using Phusion High-Fidelity DNA Polymerase (Thermo Fisher Scientific) and the primers BamHI_OSH6_for and OSH6_NotI_rev. The PCR product was digested with BamHI and NotI and ligated into the pCM189 expression vector digested with the same restriction endonucleases.

Plasmids pCM189-*OSH6Δ35*, pCM189-*OSH6Δ42*, and pCM189-*OSH6lidless* were generated by inverse PCR using pCM189-*OSH6* as the template and primer pairs (Table **S1**) designed to delete the corresponding regions of the *OSH6* coding sequence. PCR amplification was performed using Phusion High-Fidelity DNA Polymerase (Thermo Fisher Scientific) according to the manufacturer’s recommendations. The resulting linear PCR products were treated with FastDigest DpnI (Thermo Fisher Scientific) at 37 °C for 15 min to remove methylated template plasmid DNA. The DpnI-treated PCR products were assembled using the NEBuilder HiFi DNA Assembly Cloning Kit (New England Biolabs) according to the manufacturer’s instructions. Assembly reactions were incubated at 50 °C for 1 h and subsequently transformed into competent *Escherichia coli* cells supplied with the kit. Transformants were selected on LB agar plates supplemented with ampicillin and incubated overnight at 37 °C. All plasmid constructs were verified by sequencing.

## Discussion

In this work, we investigated whether cup-like LTPs exploit membrane contact sites to enhance the efficiency of lipid transport. Indeed, lipid transfer by LTPs frequently occurs at MCSs, where donor and acceptor membranes are brought into close proximity by membrane-targeting domains, dedicated tethering proteins, or both (3, 46). Osh4 and Osh6, however, can tether opposing membranes using their lipid-transfer domains alone (11). This raises the possibility that close membrane apposition does more than localize Osh proteins between donor and acceptor membranes and instead participates directly in lipid transfer.

A first indication of such a mechanism came from experiments with Osh6^lidless^. This mutant exhibits enhanced membrane association and little detectable cargo removal from donor membranes in the absence of an acceptor, yet retains efficient PI4P transport. Thus, the ability to remove PI4P from an isolated donor membrane can be uncoupled from the ability to transport PI4P between membranes. This suggests that PI4P extraction can occur while Osh6^lidless^ engages donor and acceptor membranes simultaneously, without requiring complete dissociation from the donor before engagement of the acceptor. PS behaves differently: although PI4P transport remains largely preserved in Osh6^lidless^, PS transport is markedly impaired, indicating distinct effects of lid removal on transport of the two Osh6 cargoes.

The cellular phenotypes further show that Osh6^lidless^ is not simply a non-functional protein. It rescues growth of the otherwise non-viable *osh1Δ–7Δ pMET3-OSH2* strain, demonstrating that it retains the ability to substitute for essential Osh functions *in vivo*. In contrast, Osh6^lidless^ fails to complement the *dpl1Δ osh6Δ osh7Δ psd1Δ* strain, consistent with its strongly impaired PS transport *in vitro*. Thus, complete removal of the lid selectively affects distinct Osh-dependent functions rather than globally inactivating the protein.

The unusual behavior of Osh6^lidless^ could nevertheless reflect a mutant-specific transport mechanism. Alternatively, lid removal may stabilize or exaggerate a normally short-lived state that is also sampled by wild-type Osh6. Our data support the latter interpretation. Wild-type Osh6 shows little PI4P removal from PS-rich donor membranes in the absence of an acceptor, yet efficiently removes and transfers PI4P when a suitable second membrane is present. Similarly, PS removal from negatively charged donor membranes is inefficient in isolation, whereas PS transport remains efficient when the acceptor membrane is also negatively charged. Thus, for both cargoes, wild-type Osh6 can efficiently transfer lipid under conditions in which removal from the isolated donor is strongly disfavored. The membrane-bridged state emphasized by Osh6^lidless^ therefore appears to be accessible also to the wild-type protein under appropriate membrane conditions.

Membrane tethering provides a physical basis for such a state. Tethering by wild-type Osh6 was strongly dependent on cargo and membrane composition. PI4P consistently suppressed detectable tethering, whereas pronounced tethering was maintained under conditions supporting efficient PS transport between negatively charged membranes. Moreover, the non-transferable PS analogue phPS supported tethering, while incorporation of PI4P into the same phPS-containing membrane abolished detectable tethering. Cargo recognition alone is therefore insufficient to destabilize the tethered state. Instead, the loss of tethering upon incorporation of PI4P into phPS-containing membranes suggests that productive PI4P extraction promotes destabilization of the membrane-bridged state.

We therefore propose a “kiss-and-ride” mechanism in which transient membrane contact facilitates PI4P extraction—the “kiss”. Productive PI4P extraction then destabilizes the bridged state, allowing PI4P-loaded Osh6 to dissociate and continue the transport cycle—the “ride”. In this model, the lipid-transfer domain transiently spans the confined space between two closely apposed membranes, allowing the opposing membrane to facilitate cargo extraction without forming a continuous lipid-conducting conduit. The lifetime of this bridged state appears to depend on cargo identity and membrane composition: whereas PI4P destabilizes detectable membrane tethering, pronounced tethering persists under conditions supporting efficient PS transport between negatively charged membranes. Membrane bridging may therefore constitute a dynamic transport intermediate whose lifetime changes during the lipid-transfer cycle.

Such a mechanism may be particularly relevant to PS/PI4P exchange at ER–PM contact sites. At the plasma membrane, PI4P is embedded in a membrane containing a large excess of PS, a condition under which PI4P extraction by Osh6 strongly depends on the presence and composition of an opposing membrane. Although the ER membrane is generally relatively neutral, its lipid distribution and surface charge may be locally heterogeneous (47), particularly at contact sites organized by Ist2. Ist2, an adaptor for Osh6, has also been proposed to function as a PS scramblase (10, 12), translocating newly synthesized PS from the luminal to the cytosolic ER leaflet (48). Local PS enrichment could therefore generate a negatively charged cytosolic ER surface opposing the negatively charged plasma membrane, creating conditions favorable for simultaneous Osh6 interaction with both membranes. Osh6 could transiently bridge the ER and plasma membranes during PS transfer, whereas subsequent acquisition of PI4P at the plasma membrane would destabilize the bridged state and allow PI4P-loaded Osh6 to return toward the ER.

The kinetic measurements provide independent line of support for this model. Depending on membrane composition, lipid transport was either substantially slower than cargo release from preloaded Osh6 or approached the cargo release rate. Importantly, the latter behavior occurred under conditions in which cargo removal from the isolated donor membrane was inefficient. Thus, cargo acquisition from an isolated donor is strongly disfavored under these conditions, yet a suitable opposing membrane allows transport to proceed at a rate approaching cargo release from preloaded Osh6. Together with enhanced membrane tethering under the same membrane conditions, this behavior supports a model in which engagement of the opposing membrane facilitates cargo extraction within a transient membrane-bridged state.

Although Osh6 lacks extended intrinsically disordered regions (IDRs), its activity at ER–PM contact sites is coupled to Ist2, which contributes to maintaining close membrane apposition. More generally, many MCS proteins, including members of the ORP/Osh family, contain extended IDRs that can influence intermembrane spacing and contact-site organization (49–51). Formation of the membrane-bridged intermediate proposed here may therefore depend not only on the membrane-binding properties of the lipid-transfer domain but also on the geometry imposed by the surrounding MCS machinery. In Osh6, Ist2 may create an intermembrane environment favorable for transient engagement with both membranes, whereas multidomain ORPs may impose analogous geometric constraints through their own membrane-targeting and disordered regions. MCS geometry may therefore regulate the balance between shuttle-based and membrane-bridged transport.

Together, our results suggest that transient membrane bridging is an integral component of lipid transport by cup-like LTPs, directly facilitating cargo extraction and coupling lipid transfer to membrane contact dynamics. More broadly, membrane contact sites may not simply bring lipid-transfer proteins and their target membranes together, but actively shape the mechanism by which lipids are transported between them.

## Acknowledgement

We gratefully acknowledge C. Beh (Simon Fraser University, Canada) and C. Loewen (University of British Columbia, Canada) for providing the yeast strains used in this study. We thank Tamas Balla for critically reading the manuscript and providing valuable comments.

## Author contributions

AB, AE, RH, and JH performed the experiments, AB, RH and JH evaluated the data, PG, EB, AB and JH wrote the manuscript.

