## Supplementary material for "A Kiss-and-Ride Mechanism for Lipid Exchange at Membrane Contact Sites": Supplementary Information.pdf

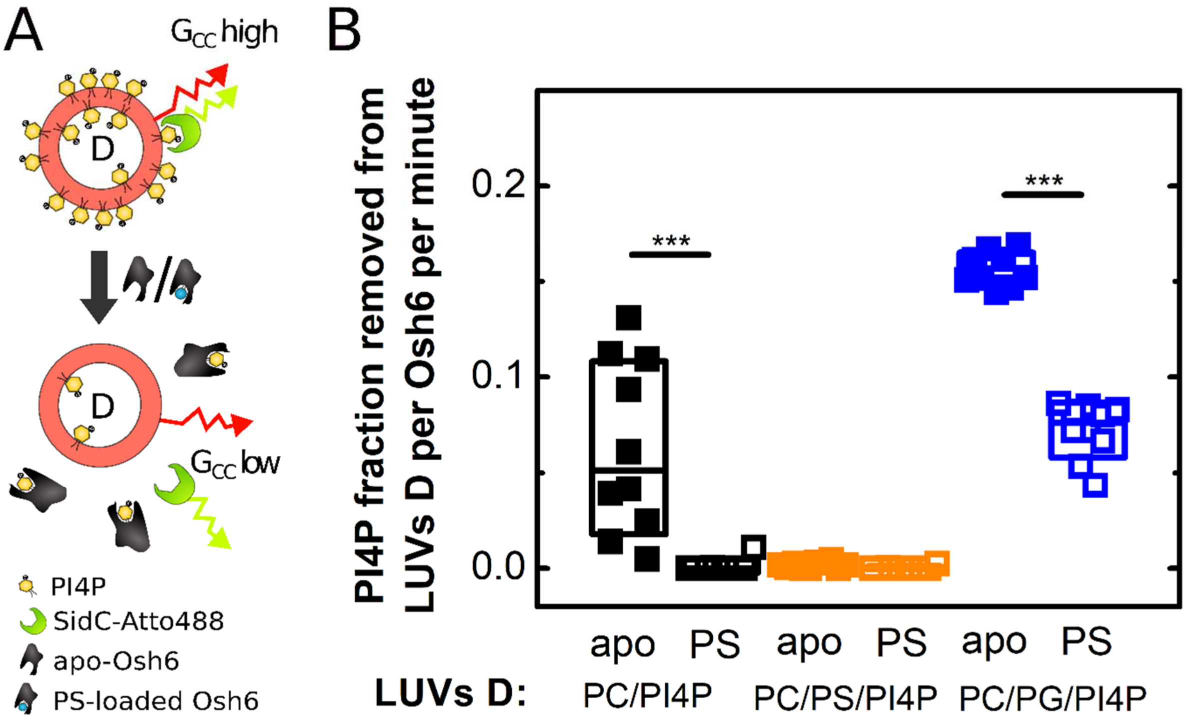


**Figure S1. Effect of PS loading of Osh6 on PI4P removal from donor membranes.**

A) Scheme of the PI4P removal assay using apo or PS-loaded wild-type Osh6. Apo Osh6 or Osh6 preloaded with PS was added to donor LUVs in the absence of acceptor LUVs, and PI4P removal was monitored by FCCS.

B) PI4P removal by apo (solid symbols) and PS-loaded (open symbols) wild-type Osh6. Donor LUVs were composed of POPC/PI4P (1 mol%) (black), POPC/PS (19 mol%)/PI4P (1 mol%) (orange), or POPC/PG (19 mol%)/PI4P (1 mol%) (blue).

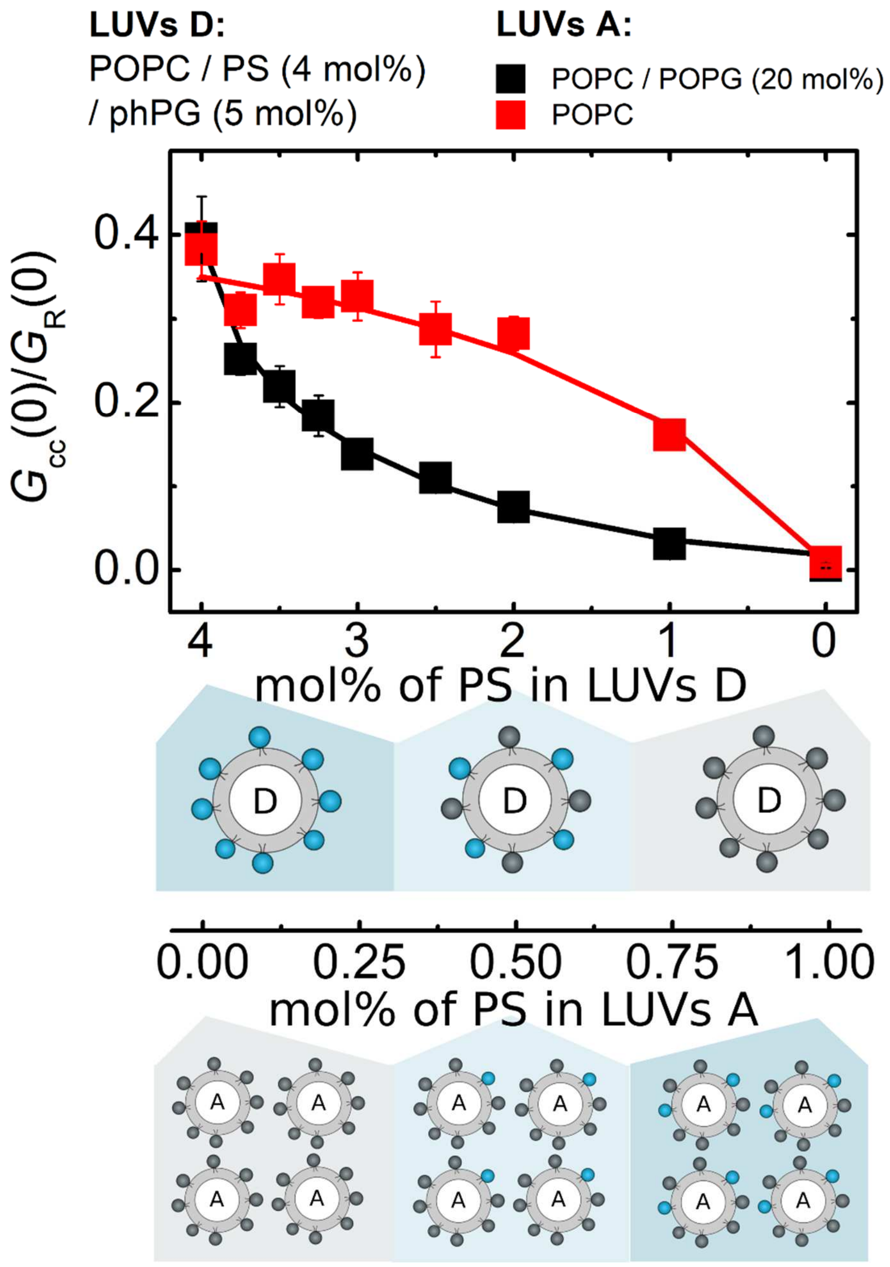


**Figure S2. Calibration of the FCCS response for PS transport.**

Mixtures of donor (LUVs D) and acceptor (LUVs A) vesicles (1:4) were prepared to represent defined extents of PS transport by progressively decreasing the PS content in LUVs D and increasing it in LUVs A. The normalized FCCS parameter, $G_{cc}(0)/G_R(0)$, was measured for each mixture and plotted as a function of the extent of PS transport. Apart from the varying PS content, donor LUVs contained POPC/1,2-diphytanoyl-sn-glycero-3-phospho-(1'-rac-glycerol) (phPG 5 mol%). PhPG was included to provide the negative membrane charge required for proper biosensor function. Acceptor LUVs were either neutral, composed of POPC (red squares), or negatively charged, composed of POPC/PG (20 mol%) (black squares).

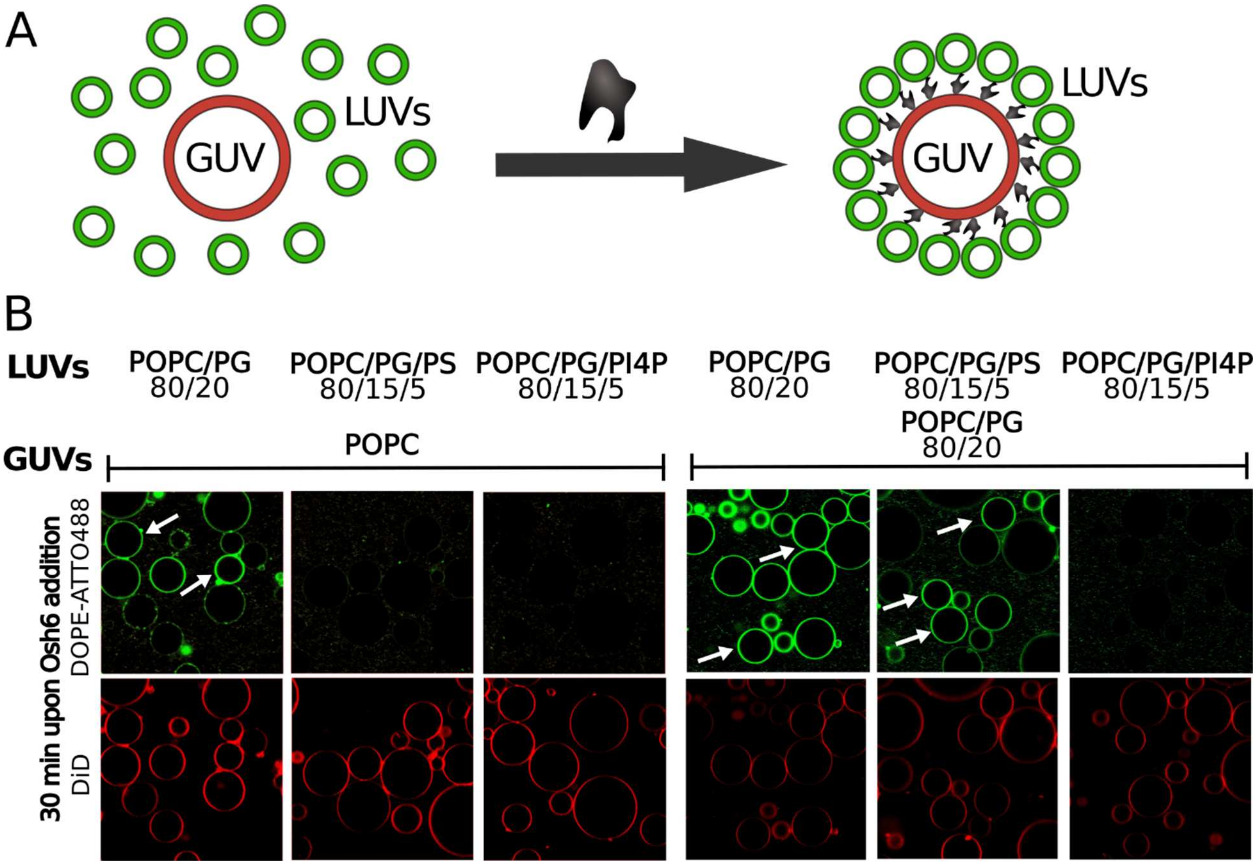


**Figure S3. Tethering of neutral and negatively charged GUVs with LUVs of different lipid compositions.**

A) Scheme of the LUV–GUV tethering assay. Osh6-mediated tethering recruits DOPE-ATTO488-labeled LUVs (green) to DiD-labeled GUVs (red).

B) Visualization of membrane tethering mediated by wild-type Osh6. DOPE-ATTO488-labeled LUVs of the indicated lipid compositions were incubated with DiD-labeled GUVs for 30 min in the presence of Osh6. GUVs were composed of either POPC or POPC/PG (20 mol%). LUVs were composed of POPC, POPC/PG (20 mol%), POPC/PG (15 mol%)/PS (5 mol%), or POPC/PG (15 mol%)/PI4P (5 mol%). White arrows indicate GUVs decorated with tethered LUVs. Total lipid concentrations were 100 µM for LUVs and 12.5 µM for GUVs; Osh6 concentration was 250 nM.

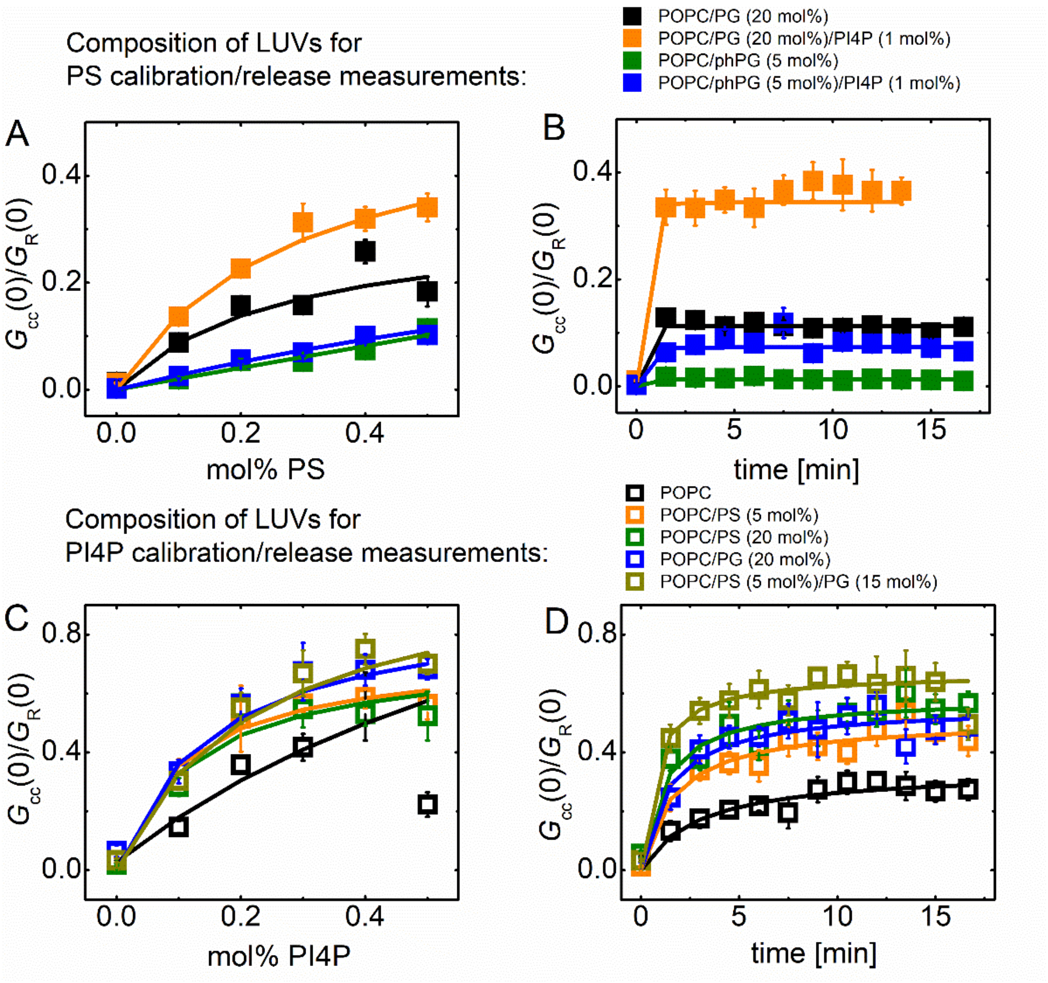


**Figure S4. Calibration and measurement of PS and PI4P partitioning between Osh6 and LUVs.**

A) Calibration of the FCCS response to the PS content of LUVs of different compositions.

B) Time-dependent increase in the FCCS signal following addition of PS-loaded Osh6 to cargo-free LUVs of the corresponding compositions. LUV compositions in A and B (solid symbols) were POPC/PG (20 mol%) (black), POPC/PG (20 mol%)/PI4P (1 mol%) (orange), POPC/phPG (5 mol%) (green), and POPC/phPG (5 mol%)/PI4P (1 mol%) (blue).

C) Calibration of the FCCS response to the PI4P content of LUVs of different compositions.

D) Time-dependent increase in the FCCS signal following addition of PI4P-loaded Osh6 to cargo-free LUVs of the corresponding compositions. LUV compositions in C and D (open symbols) were POPC (black), POPC/PS (5 mol%) (orange), POPC/PS (20 mol%) (green), POPC/PG (20 mol%) (blue), and POPC/PS (5 mol%)/PG (15 mol%) (yellow).

| Name | Sequence | Description |
|---|---|---|
| BamHI_OSH6_for | ATTGGATCCATGGGCTCCAAAAAACTGACCGTA | Forward primer to construct pCM189-*OSH6* |
| OSH6_NotI_rev | ATTGCGGCCGCCTATTGTTTTGCTGGGTTCTGCT | Reverse primer to construct pCM189-*OSH6* |
| Osh6D35_F1 | GATACCGACGATATTGATGAAGACG | Forward primer to construct pCM189-*OSH6*$^{Δ35}$ |
| Osh6D35_R1 | TTCATCAATATCGTCGGTATCCATGGATCCCCCGAATTGATC | Reverse primer to construct pCM189-*OSH6*$^{Δ35}$ |
| Osh6D42_F1 | GACGATGAATCTGGTCATAATATTATCTTG | Forward primer to construct pCM189-*OSH6*$^{Δ42}$ |
| Osh6D42_R1 | ATTATGACCAGATTCATCGTCCATGGATCCCCCGAATTGATC | Reverse primer to construct pCM189-*OSH6*$^{Δ42}$ |
| Osh6D69_F1 | CCTACTTTTATTTTAGAAAAAAAATCGATGC | Forward primer to construct pCM189-*OSH6*$^{lidless}$ |
| Osh6D69_R1 | GATTTTTTTTCTAAAATAAAAGTAGGCATGGATCCCCCGAATTGATC | Reverse primer to construct pCM189-*OSH6*$^{lidless}$ |

**Table S1.** List of primers.
